# Nonphotochemical quenching changes with abiotic stressor and developmental stages

**DOI:** 10.1101/2025.05.14.654125

**Authors:** Seema Sahay, Gabriel de Bernardeaux, Diana Gamba, Jesse R. Lasky, Katarzyna Glowacka

**Affiliations:** Department of Biochemistry and Center for Plant Science Innovation, University of Nebraska-Lincoln, Lincoln, NE, USA; Department of Biology, Pennsylvania State University, University Park, PA 16802, USA; Institute of Plant Genetics, Polish Academy of Sciences, 60-479 Poznań, Poland

**Keywords:** Nonphotochemical quenching kinetics, abiotic stresses, crops, fluorescence imaging approaches, drought, chilling, low nitrogen stress

## Abstract

Nonphotochemical quenching (NPQ) is a critical photoprotective mechanism in plants, safeguarding photosystem II (PSII) and PSI from photodamage under abiotic stress. However, it is unclear if different stressors lead to similar NPQ phenotypes, and the magnitude of natural variation (between and within plant species) in NPQ response to abiotic stress is unknown. Testing a semi-high-throughput leaf-disc approach for examining the NPQ kinetics parameters, we investigated NPQ under chilling, drought and low nitrogen stress across multiple species and/or genotypes. Our results show substantial variation in NPQ phenotypes across species, genotypes and treatments. In C3 crops, tobacco and soybean, multiple NPQ parameters generally increased under chilling and drought, while in C4 crops, maize and sorghum, NPQ traits were more variable including a decrease of multiple NPQ parameters. Low-N stress revealed genotype- and developmental stage-specific effects on NPQ, potentially reflecting distinct adaptive strategies and regulatory changes in NPQ stress response. A significant effect of ecotype and stress treatment was detected on most NPQ kinetics traits in *Arabidopsis thaliana*, however, the interaction between ecotype and treatment was stronger in drought than in chilling. Differential regulation of NPQ could be associated with a combination of changes in proton motive, ATPase synthase activity, and PSI redox state. Our findings highlight that interpreting relative changes in NPQ under abiotic stress is inherently complex and demands a broader integration of physiological data across multiple regulatory layers.

## Introduction

Photosynthesis requires plants to balance light energy capture with its use for CO assimilation (Kaiser et al., 2018; Long et al., 2022). Excess light energy, if unregulated, leads to reactive oxygen species (ROS) generation, photodamage, and reduced photosynthetic efficiency. Among photoprotective responses, one of the fastest is releasing the excess absorbed energy as heat which is observable as a phenomenon named nonphotochemical quenching of chlorophyll fluorescence (NPQ) (Müller et al., 2001; Ruban et al., 2007; Nicol et al., 2019). For NPQ to happen the antenna complexes undergo conformational changes, stimulated by an increase in the proton gradient across the thylakoid membrane, protonation of photosystem II subunit S (PsbS) and conversion of xanthophyll cycle pigments to zeaxanthin by violaxanthin de-epoxidase (VDE) (Li et al., 2000). In low light or dark, the zeaxanthin is reversed back to violaxanthin by zeaxanthin epoxidase (ZEP). The mechanisms described above contribute to two main components of NPQ, energy-dependent quenching (qE, relaxing within a minute) and zeaxanthin-dependent quenching (qZ, exhibits a slower relaxation period lasting typically 10 to 15 minutes) (Kress and Jahns, 2017). In natural conditions plants are subjected to constant changes in light intensity over time scales of seconds. These changes alter the quantity of light energy available to be captured by photosynthetic pigments requiring faster NPQ for plant fitness under fluctuating light (Burgess et al., 2016; Kaiser et al., 2018). Delayed NPQ induction or recovery can cause photodamage, while excessive NPQ reduces photosynthetic efficiency (Frenkel et al., 2007; Davis et al., 2017; Kanazawa et al., 2020). Faster NPQ adjustments has been linked to greater photosynthetic capacity and biomass in tobacco, soybean, and rice (Kromdijk et al., 2016; De Souza et al., 2022; Xin et al., 2023).

Abiotic stresses such as drought, chilling, and low nitrogen disrupt energy balance and increase the risks of photodamage (Zgallaï et al., 2005; Tantray et al., 2020; Guo et al., 2021). For instance, chilling stress reduces enzymatic rates which limits the sinks for the absorbed excitation energy that can lead to the extensive absorption of energy and ROS production. Despite the importance of NPQ in minimizing stress-induced damage, comparative studies across stresses and natural genetic variation in crops are scare (Malnoë, 2018).

NPQ as a fluorescence-based trait can be cheaply and noninvasively measured using a fluorescence imager or fluorometer (Baker, 2008). Traditional NPQ measurements are low-throughput, limiting their utility in large-scale studies. Surveying variation in NPQ kinetics across large plant populations in natural or field conditions is highly challenging. Recently a semi-high-throughput approach has enabled simultaneous NPQ measurement across many genotypes, improving feasibility under field conditions (Sahay et al., 2023). This approach involves the use of small leaf sections, punched from plants, which can be fit into a 96-well plate to analyze simultaneously the large number of plants using the fluorescence imaging system. This approach enables the collection of a time series of points that can be used in NPQ kinetics studies. The leaf-disc approach offers the added benefit of overnight adaptation, which helps minimize the impact of microenvironmental conditions such as light intensity, temperature, and time of day during sampling. Lately, the application of this approach has identified novel NPQ regulatory genes, including *Atypical Cys His Rich Thioredoxin 3*, *Chloroplast Outer Envelope Protein 37*, *Plastid Movement Impaired 1*, *Thioredoxin Y1*, *Non-Yellowing 1, L-galactono-1,4-lactone dehydrogenase* or minor PSII antenna protein *CP24* (Sahay et al., 2023; 2024a; Ferguson et al., 2025; Kumari et al 2025). However, it has to be still determined if this approach works for various plant species grown under different stress conditions.

In this study, we investigated NPQ in four crops, tobacco (*Nicotiana tabacum*), soybean (*Glycine max*), maize (*Zea mays*), and sorghum (*Sorghum bicolor*), under chilling and drought stresses. We also assessed natural variation in NPQ traits in maize, teosinte (*Z. mays* subsp. *parviglumis*) and *Arabidopsis thaliana* collectively in three abiotic stresses. By comparing the leaf-disc and whole-plant level sampling approaches, we revealed that leaf-disc fluorescence measurements are highly reliable in studying NPQ kinetics under control and stress conditions. We hypothesized that different abiotic stresses would lead to the same NPQ phenotype of faster and stronger NPQ induction and higher NPQ residuals which should minimize oxidative damage. However, NPQ traits across multiple species and/or genotypes showed no uniform NPQ phenotype under chilling, drought and low nitrogen. While changes in NPQ generally followed the change in proton motive forces, only in coupled species × stress combinations could they be tied to changes in the regulation of chloroplast ATP synthase. Comparison of NPQ under the low soil fertility in different development stages in a set of maize genotypes revealed a significant effect of both plant ontogenesis and genotype on NPQ kinetics. Similarly, a significant effect of ecotype and treatment was detected on most NPQ kinetics traits in *Arabidopsis thaliana*, however, the interaction between ecotype and treatment was stronger in drought than in chilling.

## Results

### Comparison of leaf-disc and whole-plant measurement approach of NPQ for control- and stress-condition-grown plants

We tested the leaf-disc- and whole-plant-based approach to measure NPQ kinetics across various plant species and stress conditions using Pulse-Amplitude-Modulation (PAM) fluorometry. First, we tested plants grown under control conditions that offer broad NPQ variation by including *PsbS*-silencing (psbs-4; exhibiting low NPQ; Glowacka et al., 2018), *PsbS*-overexpressing (PSBS-28; exhibiting high NPQ; Głowacka et al., 2018) lines and corresponding wild type (WT) of tobacco. Under control conditions, NPQ induction and relaxation kinetics measured in leaf discs were very similar to measurements performed in the whole plant for all three investigated lines (Fig. S1A-C). A very close to one-to-one relationship was observed between NPQ values at each time points derived from leaf-disc and whole-plant approaches when linear regression was inspected for WT (*y* = 0.97*x* - 0.047; R^2^ = 0.99, *P* < 0.0001, Fig. 1A), psbs-4 (*y* = 1.02*x* - 0.039; R^2^ = 0.99, *P* < 0.0001, Fig. 1B), and PSBS-28 (*y* = 0.96*x* - 0.076; R^2^ = 0.99, *P* < 0.0001, Fig. 1C). The NPQ parameters obtained from the hyperbolic fits of the NPQ kinetics curve under light and dark conditions were not significantly different between both approaches despite huge variation in NPQ phenotype between three tested lines (*P* ≥ 0.06; Fig. S1 D-I; Table S1). The difference between measurement approaches for all parameters, on average, showed insignificant differences of 6.2% (from 0.4% to 14%; Fig. S1D-I).

**Figure 1.**
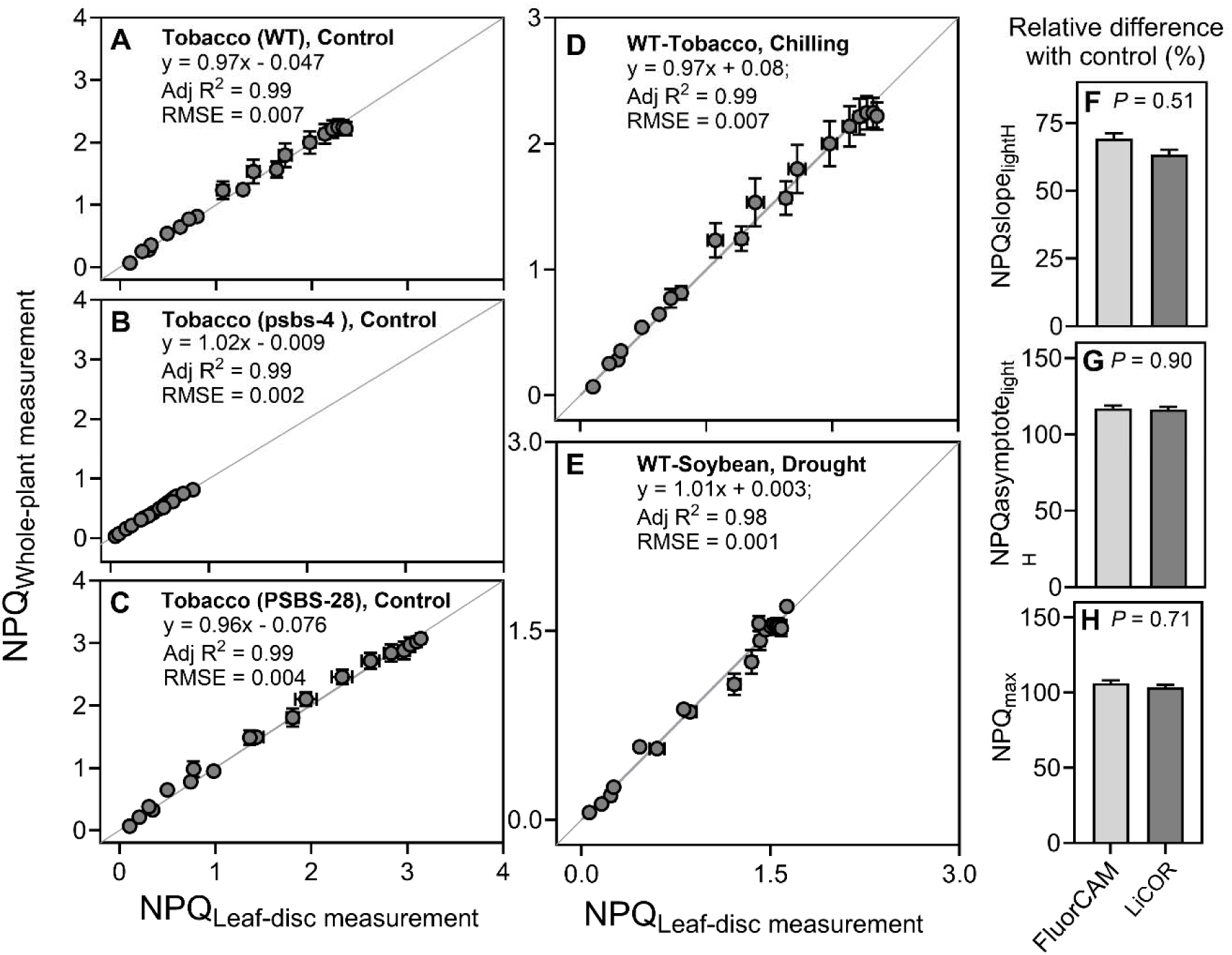
Comparison of approaches used for NPQ measurements. (**A-E)** Comparison of leaf-disc- and whole-plant-based approach for NPQ measurement under control and stress conditions. (A-C) linear relationship between leaf disk and whole plant measurements of NPQ induction and relaxation kinetics in wild type (WT) and two transgenics (psbs-4 silencing line and PSBS-28 overexpressing line) of tobacco (*Nicotiana tabacum*) plants grown under control conditions. (**D, E)** linear relationship between leaf-disc measurement and whole-plant measurement of NPQ induction and relaxation kinetics in WT tobacco and soybean (*Glycine max*) plants grown under chilling and drought, (**F-H)** Comparison of NPQ traits collected using FluorCAM fluorescence imager or LiCOR fluorometer with a multiphase flash routine base on the relative change under chilling in three NPQ induction traits in sorghum plants (*Sorghum bicolor*). In panel F-H, the values of traits in chilling plants were expressed as relative changes to corresponding measurements in plants grown under control conditions. NPQslope_lightH_ - rate of NPQ induction; NPQasymptote_lightH_ - steady-state of NPQ induction and NPQ_max_ – maximum value of NPQ induction (a full definition of each trait is provided in Table S1). Adj R^2^ - coefficient of determination for linear relation; RMSE - root mean square error. In panels A-C, D data represent the mean ± SEM (tobacco, n = 8; soybean, n = from 4 to 5 biological replicates, each from two to three technical replicates). In panels F-H, data represent the mean ± SEM (sorghum, n = 9). In panels F-G, the *P*-value represents the effect of the method in the ANOVA. Data present results of Experiment I.

Next, we compared results from leaf disc and whole plant measurements on plants experiencing abiotic stress to find out if the linear relation between both sets of data still holds. For both chilling-treated tobacco and drought-treated soybean wild type plants, NPQ induction and relaxation kinetics measured in leaf discs were very similar to measurements performed in whole plants (Fig. S2 A-B). Leaf discs and whole plant NPQ data showed strong linear relation under both stresses (Fig. 1D, E; y = 0.97x + 0.08; R^2^ = 0.99 in chilling and y = 1.01x - 0.003; R^2^ = 0.98 in drought). Furthermore, the NPQ traits delivered from hyperbolic fitting to NPQ kinetics curves were statistically no different between both approaches (*P* ≥ 0.08; Fig S2 C-J). The exception was NPQslope_lightH_ and NPQslope_darkH_ in chilling treated tobacco which showed significant differences in values obtained from both approaches (*P* = 0.03 and *P* = 0.001; Fig S2 C, D) with a lack of consistency in directionality of differences between them. While both NPQ measurement approaches displayed reliable data distribution allowing for the fitting of light and dark points to hyperbolic equations with high significance (goodness of fit from 0.95 to 0.99), the leaf-disc approach exhibited either similar (tobacco NPQgof_dark_; *P* = 0.3) or significantly higher goodness of fit compared to the whole-plant approach for NPQ data (NPQgof_dark_ soybean and NPQgof_light_ tobacco and soybean; *P* ≤ 0.03; Fig. S2 I, J). Furthermore, across control, chilling, and drought conditions in tobacco and soybean plants, maximum quantum yield of photosystem II (*F*_v_/*F*_m_), as an estimation of the loss of function of photosystem II (PSII) reaction centers in response to stresses were not significantly different between leaf-discs and whole-plant measurements (the differences were from 0.0 to 2.4%; *P* ≥ 0.05; Table S2). In addition, we also compared data collected from leaf discs using FluroCam fluorescence imager with data collected on the excised fully expanded leaves using a multiphase flash routine in LI-6800 fluorometer (LI-COR, Inc. Lincoln, NE, USA). For this approach we used sorghum plants under control and chilling stress conditions and measured NPQ kinetics in light. While absolute values of NPQ differed between both instruments, the relative changes between chilling stress and control conditions were very similar (Fig. 1). The relative changes with control differed on average by 3.9% between data obtained with the use of FluroCam-imager and Li-COR-fluorometer for all three NPQ traits obtained from hyperbolic fitting (*P* ≥ 0.51; Fig. 1F-H; Table S3).

### Effect of different abiotic stresses on NPQ kinetics in various crops

To explore changes in NPQ kinetics and its traits under chilling and drought stress various economically important crops were investigated. Between these crops, both C_3_ (tobacco and soybean) and C_4_ (maize and sorghum) photosynthesis types were represented. The tested stresses substantially affected the kinetics of NPQ, however, stress-specific differences were also observed in some species (Fig. 2).

**Figure 2.**
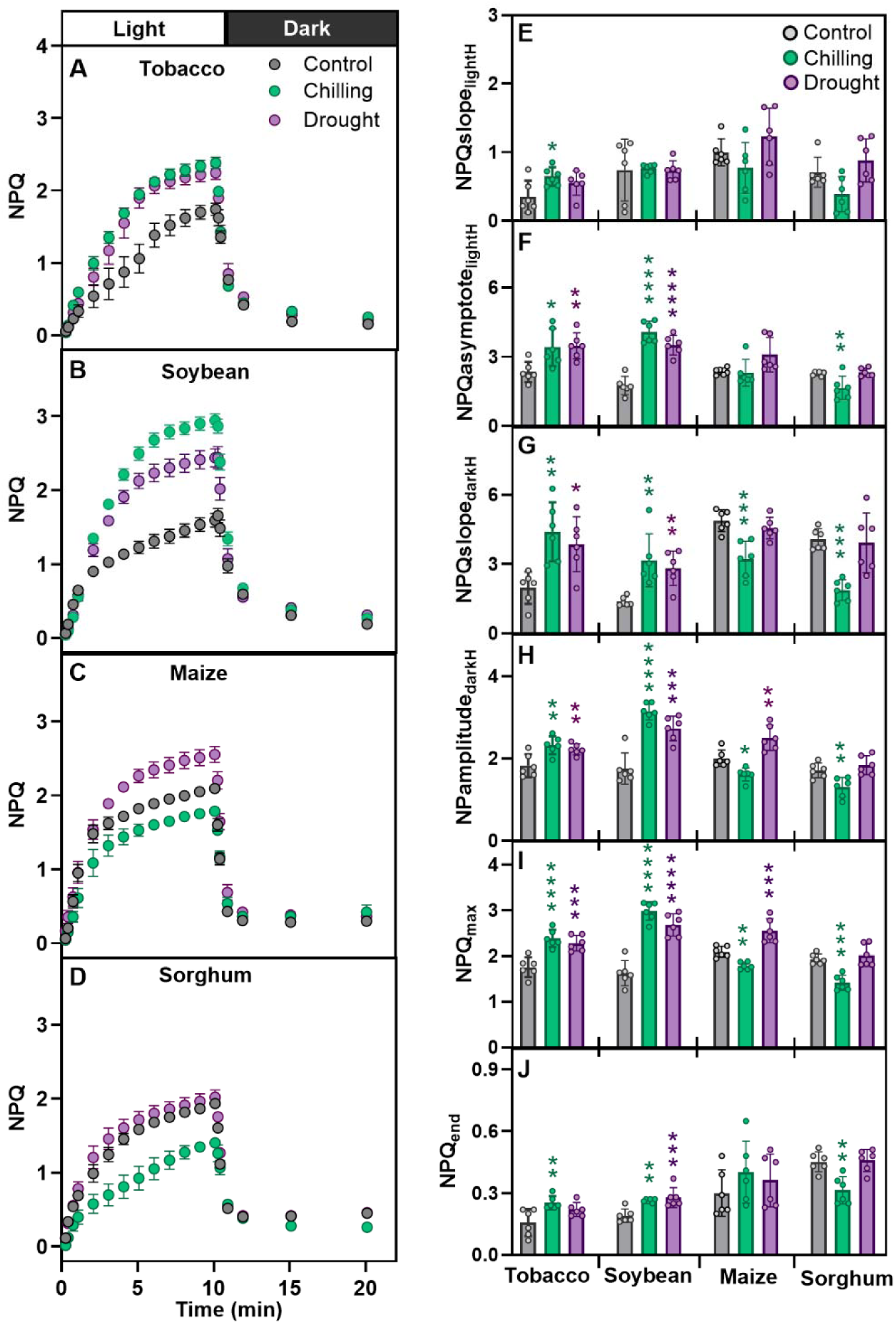
Response of NPQ kinetics across two stresses in four crops. (**A-F)** Kinetics of NPQ induction in the light (indicated by the white horizontal bar) and relaxation in the dark (indicated by the black horizontal bar) across chilling and drought stress in tobacco (*Nicotiana tabacum*), soybean (*Glycine max*), maize (*Zea mays*), and sorghum (*Sorghum bicolor*). (**G-L)** Parameters associated with NPQ kinetics in light and dark describing rate (NPQslope_lightH_), range (NPQamplitude_darkH_), steady-state (NPQasymptote_lightH_), and last value of NPQ in the light and dark (NPQ_max_ and NPQ_end_, respectively). Values are mean ± SEM (tobacco, n = 4-8; soybean, n = 4-5; maize, n = 4; and sorghum, n = 6 biological replicates, each from two to three technical replicates). In panels E-J, individual biological replicates are represented as filled circles. Presented data were collected using the leaf-disc approach. Asterisks indicate the significant differences relative to control in Dunnett’s two-way test; \**P* < 0.05; \*\**P* < 0.01; \*\*\**P* < 0.001, and \*\*\*\**P* < 0.0001). Data present results of Experiment II.

In both C3 crops both stresses significantly increased NPQ induction and relaxation parameters from 22% to 123% (*P* ≤ 0.02) for tobacco and from 39% to 133% (*P* ≤ 0.02) for soybean (Fig. 2A-D, F-I), except speed of NPQ induction (NPQslope_lightH_) and residual NPQ (NPQ_end_) (Fig. 2 E and J). Unlike in tobacco and soybean, the values of all NPQ traits decreased under chilling stress in both maize and sorghum (from -2.4% to -33.9%; Fig. 2F-I; *P* ≤ 0.02) except for residual NPQ (Fig. 2 E-J). The impact of drought on NPQ kinetics was less consistent and significant in both tested C4 crops (Fig. 2). In maize, drought significantly increased only NPQamplitude_darkH_ and NPQ_max_ by 25% and 22%, respectively (*P* ≤ 0.004) compared to other traits which showed non-significant increase or decrease (*P* ≥ 0.06; Fig. 2E-J). Conversely, in sorghum drought led to non-significant changes in all six analyzed NPQ traits.

### Effect of abiotic stresses on photosynthesis-related parameters across four crops

To elucidate the mechanism of NPQ regulation under stresses in C3 and C4 crops, simultaneously with NPQ investigation under drought and chilling stresses, we measured broadly defined photosynthetic phenotype (Fig. 3 and Fig. 4). The changes in a simplified NPQ parameter which does not require dark adaptation (NPQ_T_) measured on attached leaves (Fig. 3A-D) were largely consistent with the maximum value of NPQ estimated on leaf discs collected to plates (Fig. 2I). Tobacco and soybean showed significant increase and sorghum showed significant decrease in NPQ_T_ under both chilling and drought, while in maize chilling significantly decreased and drought significantly increased NPQ_T_ (Fig. 3A-D) (*P* ≤ 0.003). The increase of NPQ_T_ by 5-fold under chilling in soybean was the most radical changed noted among tested here species by stress combinations. The thylakoid proton motive forces (*pmf*) *in vitro* (estimated from the ECSt parameter) increased most substantially (2-fold; *P* < 0.0001) in soybean under chilling treatment, while decreased the most in maize under the same stress (−66%; *P* = 0.001; Fig. 3F, G). A significant reduction (−23%; *P* = 0.002) in *pmf* was also observed in sorghum under drought but not in chilling stress (Fig. 3H). Furthermore, in tobacco, *pmf* was increased non-significantly under both stresses (Fig. 3E). The changes in thylakoid proton conductivity, termed gH^+^, remained largely unchanged across species, with significant changes only being observed in soybean (on average 48% decrease under both stresses; *P* < 0.0001) and sorghum (51% increase under drought; Fig. 3I-L). The linear electron flow (LEF) increased significantly in tobacco (drought, *P* = 0.03) and sorghum (chilling, *P* = 0.002) (Fig. 3M-P). Oppositely, LEF was significantly reduced under chilling in soybean (*P* = 0.05) and under both stresses in maize (*P* ≤ 0.03) (Fig. 3N, M). A significant increase in LEF was observed in sorghum under chilling (*P* = 0.005; Fig. 3P). Both chilling and drought substantially decreased stomatal conductance (*gsw*) in investigated crops except for maize in drought (Fig. 3Q-T). The redox state of photosystem I (PSI) responded greatly to both stresses with significant differences (Fig. 4). Chilling and drought significantly increased the over-reduced fraction of PSI centers in tobacco and soybean (*P* = 0.002; Fig. 4E, F), which was also true for sorghum in chilling (*P* = 0.002; Fig. 4H). Both stresses only marginally affected the PSI oxidized centers in C3 crops while the significant less PSI oxidized centers were observed in maize and non-significant less in sorghum (Fig. 4C, D).

**Figure 3.**
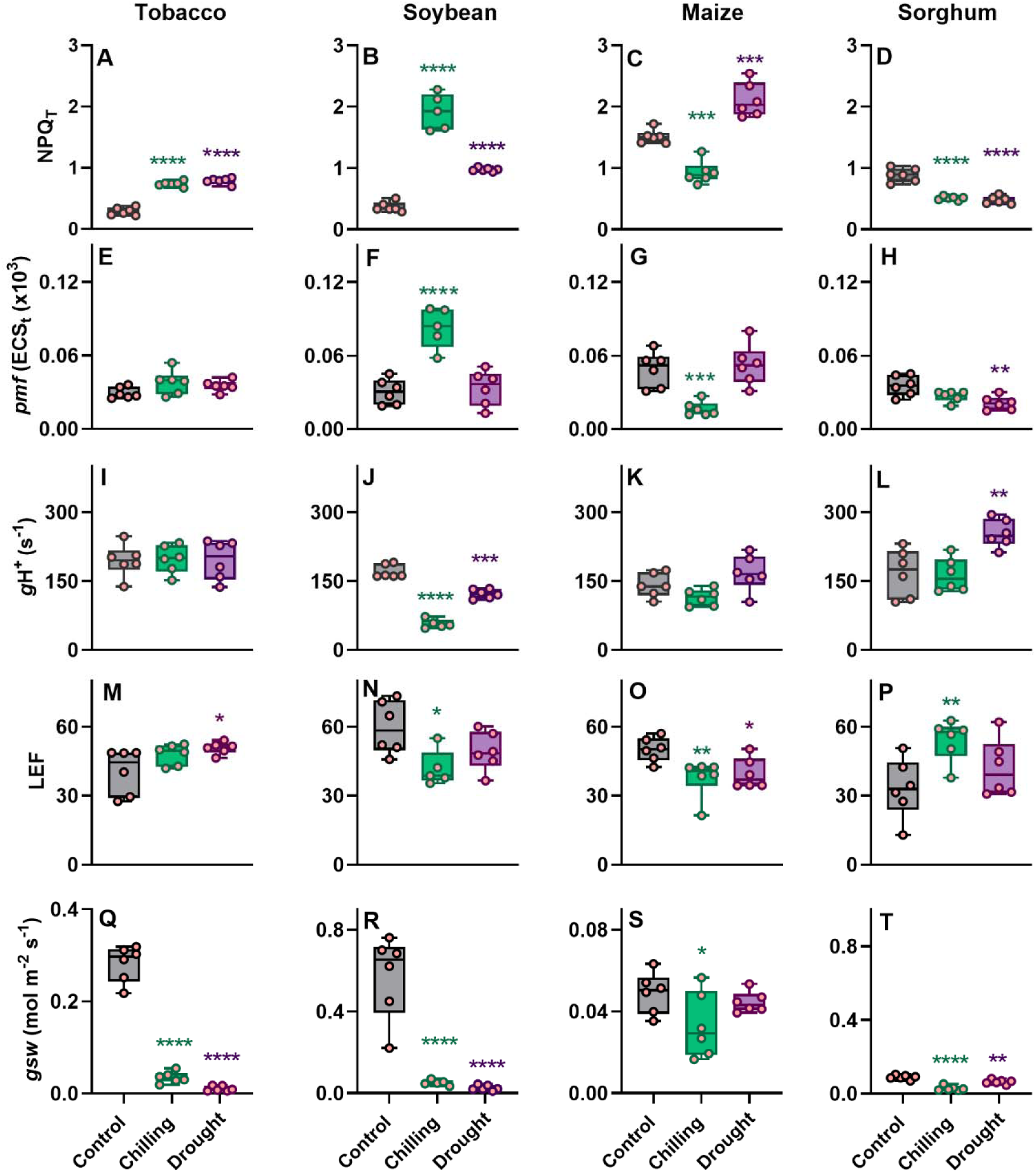
Effect of abiotic stresses on photosynthesis-related parameters across multiple crops. Theoretical NPQ (NPQ_T_), proton motive force (*pmf*; estimated as ECSt), proton conductivity of ATPase synthase (estimated as *g*H+), linear electron flow (LEF), and stomatal conductance (*gsw*), were measured in three-weeks old plants in tobacco (*Nicotiana tabacum*), soybean (*Glycine max*), maize (*Zea mays*), and sorghum (*Sorghum bicolor*) under chilling and drought stress conditions. Data represent the means ± SEM (n = 6 biological replicates, n = 5 for soybean under chilling). Asterisks indicate significant differences with control (Dunnett’s two-way test; \**P* ≤ 0.05, \*\**P* ≤ 0.01, \*\*\**P* ≤ 0.001; \*\*\*\**P* < 0.0001). Data present results of Experiment II.

**Figure 4.**
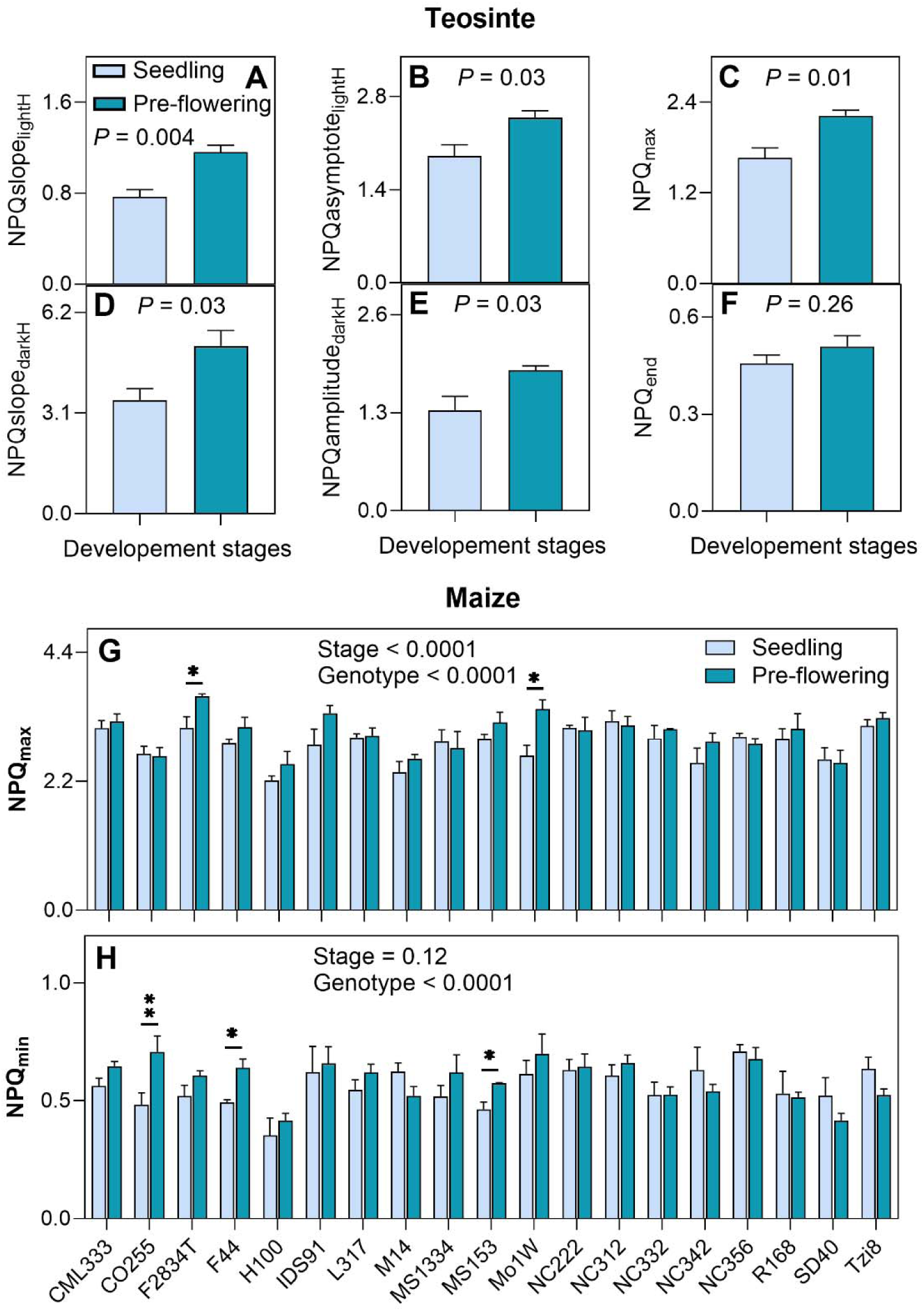
Effect of abiotic stresses on PSI redox state across multiple crops. The oxidized and over-reduced fraction of PSI centers were measured in three-weeks old plants in tobacco (*Nicotiana tabacum*), soybean (*Glycine max*), maize (*Zea mays*), and sorghum (*Sorghum bicolor*) under chilling and drought stress conditions. Data represent the means ± SEM (n = from 5 to 6 biological replicates). Asterisks indicate significant differences with control in Dunnett’s one way test; \**P* ≤ 0.05, \*\**P* ≤ 0.01, \*\*\**P* ≤ 0.001; \*\*\**P* ≤ 0.001). Data present results of Experiment II.

### Intraspecific variation in NPQ kinetics in response to a combination of fluctuating light with abiotic stresses

To evaluate intraspecific variation, the impact of fluctuating light on NPQ kinetics in response to potential high-elevation stressors, chilling (4-6°C; overnight or 1 day) and drought (5-day-holding of watering), was investigated in four *Arabidopsis thaliana* ecotypes originating from the range of elevations between 52 and 2995 m. Under fluctuating light conditions, chilling and drought stressors demonstrated distinct effects on NPQ induction and relaxation in the low and high-elevation ecotypes (ANOVA Table S4; Fig. 5; Fig. S3, S4). Under drought stress, all four ecotypes required either the same or less time for NPQ induction (smaller NPQtime_constant_light_; treatment × ecotypes; *P* = 0.06; ANOVA Table S4; Fig. S4 D-F). Oppositely, chilling-treated plants required significantly more time than control plants for NPQ induction when exposed to fluctuating light (NPQtime_constant_light_; treatment × ecotypes; *P* = 0.03; ANOVA Table S4; Fig. S3 D-F). The residual NPQ in the dark increased most substantially between 2^nd^ and 3^rd^ cycles regardless of the treatment. The low-elevation CYR ecotype, representing a seasonally dry region in France, exhibited a significantly slow NPQ response (up to 56% by 3^rd^ cycle; Fig. 5A; Fig. S3 D-F) to chilling but a significantly fast response to drought relative to control through oppositely less time needed for rapid NPQ induction (up to 228% by 3^rd^ cycle; Fig. 5B; Fig. S4 D-F). Conversely, the high-elevation Dja-1 ecotype from cold regions in Kyrgyzstan showed fairly unaltered NPQ kinetics in response to drought, yet displayed a strong response to chilling by maintaining consistently higher NPQ levels in light face during fluctuating light treatments (Fig. 5 C–D, Fig. S3, S4). In contrast, the low elevational IP-Trs-0 from mesic Iberia showed a significant increase in steady-state NPQ during 3 light-dark fluctuating light in both chilling (up to 20%) and drought (up to 13%) relative to control (Fig. 5 E–F; Fig. S3, S4 G-I).

**Figure 5.**
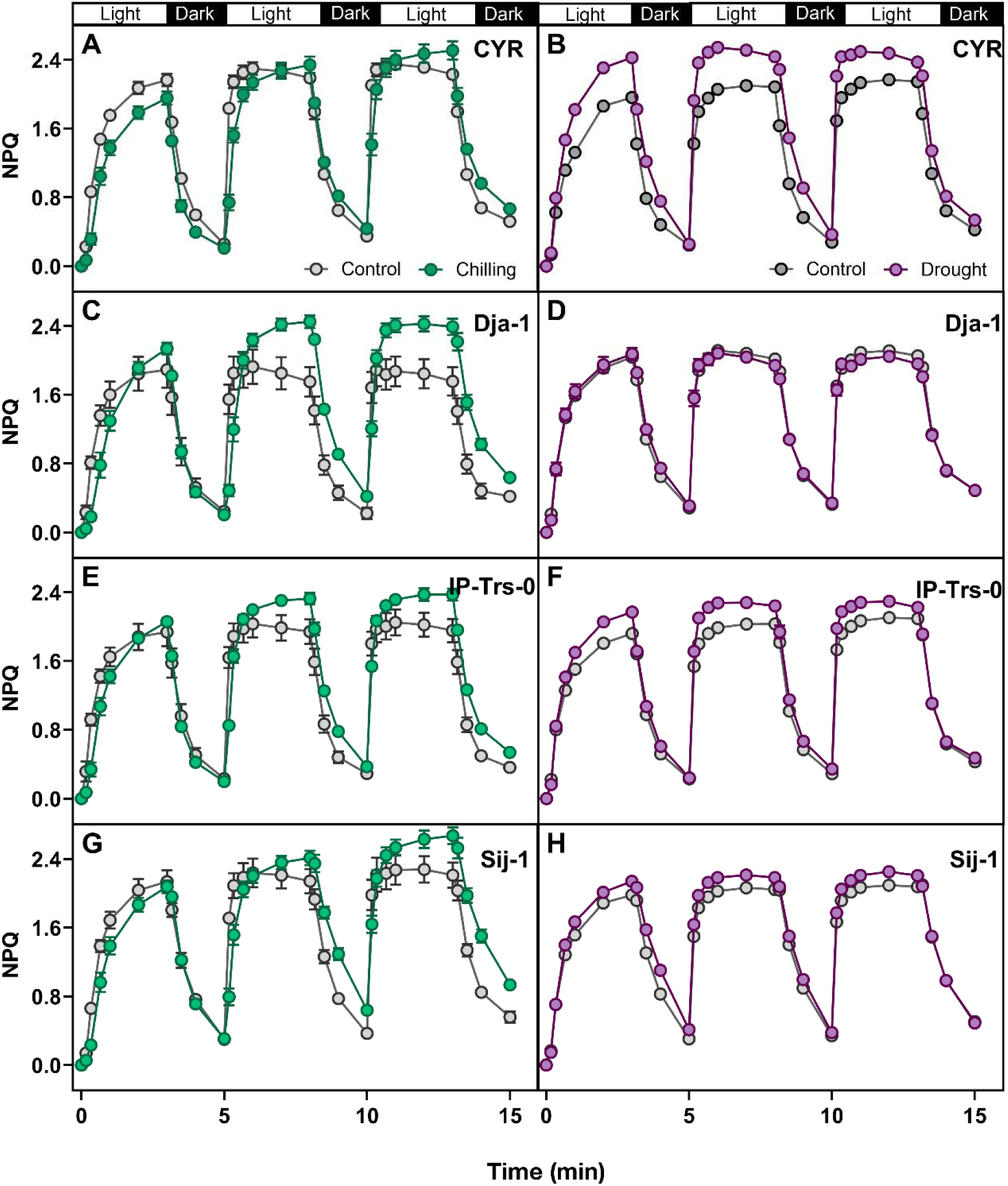
Effect of fluctuating light on NPQ in low and high elevational *Arabidopsis thaliana* ecotypes under chilling and drought conditions. Kinetics of NPQ induction and relaxation curves during three sequential light-dark cycles under cold (left panels) and drought (right panels) with corresponding control. Ecotypes represent different elevations (in the bracket): **A–B)** CYR (52 m). **C–D**) Dja-1 (2995 m). **E–F)** IP-Trs-0 (254 m). **G–H)** Sij-1 (2459 m). Presented data were collected using the whole-plant approach. Data represent the means ± SEM (n = 5-8 biological replicates). Corresponding NPQ traits and statistics are presented in supplementary data Fig. S3, Fig. S4 and Table S4. Data present results of Experiment III.

### Effect of plant developmental stage in combination with low soil fertility on NPQ kinetics

To explore the effect of the developmental stage on NPQ, we measured NPQ kinetics both at seedling and pre-flowering stages in a teosinte genotype under growth-chamber conditions and 19 maize genotypes representing inbred lines from public-sector corn-breeding programs under field conditions both at low-nitrogen treatment (Fig. 6; Fig. S5). All six investigated NPQ kinetics traits in teosinte were significantly higher at pre-flowering compared to the early-vegetative stage from 34% to 51% (*P* ≤ 0.03; Fig. 6A-E). The exception was NPQ_end_ which still showed ∼11% increase in the pre-flowering stage, however this difference was non-significant relative to the value at the early-vegetative stage (*P* = 0.26; Fig. 6F). Investigating within the set of 19 genotypes of maize, NPQ values displayed genotype-dependent variability across the two growth stages (*P* < 0.0001; ANOVA Table S5). All analyzed NPQ traits were significantly affected by the developmental stage (*P* ≤ 0.001; ANOVA), except for NPQamplitude_darkH_ and NPQ_end_ (*P* ≥ 0.10; ANOVA). Interestingly, rates of NPQ induction and relaxation were the only two traits for which significant interaction between developmental stage and genotype was detected (*P* = 0.01; ANOVA). With advancing plant age genotypes displayed non-significant changes (*P* ≥ 0.11) in NPQ_max_ except F2834T and Mo1W that showed significantly higher values at pre-flowering compared to the seedling stage (*P* ≤ 0.03; t-test; Fig. 6G). Interestingly, for NPQ_end_, three genotypes (CO255, F44 and MS153) showed significantly higher values (*P* ≤ 0.04), while the rest of the genotypes showed non-significant changes between two growth stages for NPQ_end_ (*P* ≥ 0.11; t-test; Fig. 6H). In the majority of genotypes (15 out of 19) the NPQslope_lightH_ was lower or had no change in a pre-flowering than in the early-vegetative stage with six genotypes showing a significant difference (*P* ≤ 0.05; t-test; Fig. S5A). Oppositely, NPQslope_darkH_ showed an increase in pre-flowering stage in 16 genotypes with five genotypes showing a significant difference (*P* ≤ 0.03; t-test; Fig. S5C).

**Figure 6.**
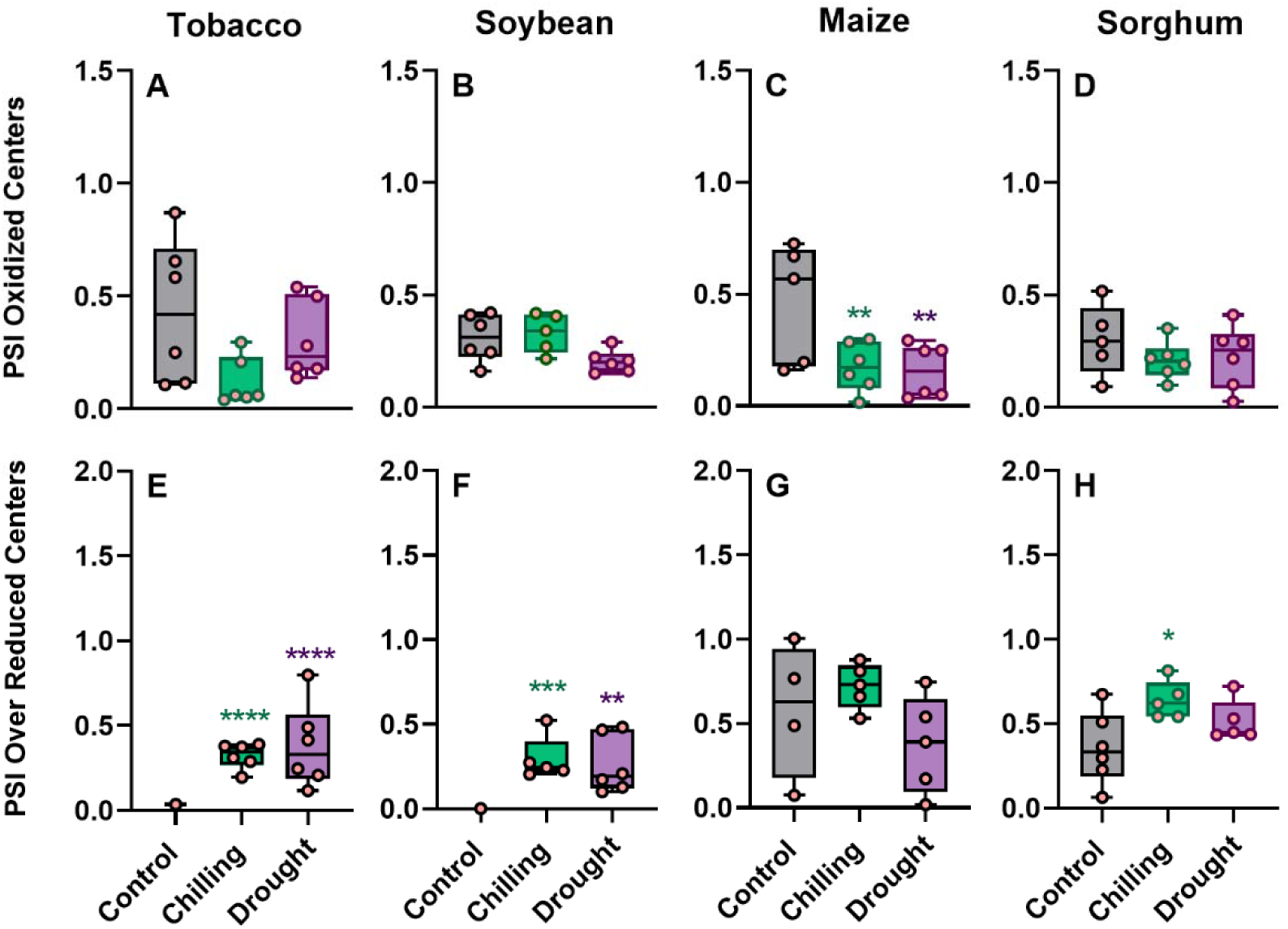
**Effect of developmental stages on NPQ kinetics**. Traits of NPQ kinetics were measured at seedling and pre-flowering stages in a teosinte (*Zea mays* subsp. *parviglumis*) line and a set of 19 maize (*Z. mays*) lines grown under growth chamber and field low N stress conditions, respectively. **A, D)** NPQslope_lightH_ and NPQslope_darkH_ describe the rate of NPQ in the light and dark, respectively. **B)** NPQasymptote_lightH_ describes the steady-state of NPQ in the light. **C, G)** NPQ_max_ is the last value of NPQ in light. **E)** NPQamplitude_darkH_ describes the range of NPQ relaxation. **F, H)** NPQ_end_ is the last value of NPQ dark. Presented data were collected using the disc-leaf approach. Data represent the means ± SEM (n = 4 biological replicates). In panels A-F, the *P*-value indicates the effect of the developmental stage in ANOVA. In panels G and H, asterisks indicate the significant differences in NPQ values between different developmental stages in t-test (\**P* < 0.05, \*\**P* < 0.01). Data present results of Experiment IV.

## Discussion

### Leaf-disc fluorescence method is reliable in NPQ kinetics studies

Our data showed that leaf-disc fluorescence measurements using a fluorescence imager are reliable for studying NPQ kinetics under control and stress conditions. Comparing data from leaf discs in 96-well plates with similar leaf areas extracted from images of intact plants showed a significant linear relationship and led to similar values of NPQ parameters (Fig. 1A-E). Furthermore, the relative changes of NPQ parameters between control and stress-treated plants were consistent. We also compared the leaf-disc approach with measurements performed on intact leaves using a multiphase flash routine applied with a fluorometer (Fig. 1F-H) (Loriaux et al., 2013). In agreement with down estimation of *F*_m_’ via fluorescence imaging due to not fully saturating pulses, the observed absolute values of NPQ parameters delivered with fluorescence imager showed the tendency to be higher than with multiphase flash routine (Table S3). In addition, other factors might contribute to observed differences between both methods coming from used instrumentation; FluorCam FC 800-C (PSI) versus LI-6800 (Li-COR), such as differences in the geometry of light source or distance of light from leaf tissue. Regarding the relative difference between genotypes and treatments, the leaf-disc fluorescence imaging approach offers a reliable, semi-high-throughput method for studying NPQ kinetics and a processing large number of samples in a short time. Together with high-density genetic marker information available for an increasing number of species leaf-disc approach for studying NPQ offers an opportunity to reveal NPQ kinetics regulatory genes via GWAS or QTL mapping as it has been recently demonstrated in multiple species (Ferguson et al., 2025; Sahay et al., 2023; 2024a; Kumari et al., 2025).

### NPQ varies across stresses, species and accessions

Our study showed no consistent NPQ phenotype under chilling, drought and low nitrogen across multiple accessions (Fig. 2, 3, 5, 6). While chilling mostly increased NPQ traits in tobacco and soybean, it decreased most of NPQ traits in maize and sorghum (Fig. 2). Increase in the initial slope of NPQ in chilling was only observed in tobacco while higher residual NPQ was observed in both tobacco and soybean but not maize or sorghum. Faster induction of NPQ would protect from chilling bright mornings and higher residual NPQ could offer more sustainable protection against stresses. While maize and sorghum are known for their low chilling tolerance (Glowacka et al., 2014), the exceptional chilling tolerance of other warm season C4 grasses, *Miscanthus*, might be due to faster NPQ induction and retention of NPQ under low temperatures (Haupt and Glowacka, 2024; Kumari et al., 2025).

Under chilling conditions, four Arabidopsis ecotypes representing contrasting climates of origin showed reduced NPQ induction speed (Fig. S3) but increased residual NPQ at the end of the 3^rd^ light-dark cycle, with no consistent change in other traits. We found a lack of a faster induction rate of NPQ in Arabidopsis accessions from the high mountains of Uzbekistan and Kyrgyzstan, Sij-1 and Dja-1. This result may suggest that some long-term photoprotective responses including anthocyanin production and a decrease in the size of light-harvesting antennae can interplay with observed NPQ phenotype (Dalal and Tripathy, 2018; Borisova-Mubarakshina et al., 2020; Haupt and Glowacka 2024) to contribute to local adaptation to high elevation (Gamba et al. 2024). Previous reports suggest also that the timing of NPQ response to stresses might differ between accessions making further complications in the direct comparison of different accessions (Riva-Roveda et al., 2016; Cackett et al., 2023; Mai et al., 2009).

Similarly to chilling, drought significantly upregulated the majority of NPQ kinetics traits in tobacco and soybean (Fig. 2). The exception was the speed of NPQ induction (NPQslope_lightH_) which was not significantly affected both in tobacco and soybean, and residual NPQ (NPQend) which was significantly affected only in soybean. In maize, only NPQamplitude_DarkH_ and NPQ_max_ significantly increased under drought, while no change was observed in all six NPQ traits in sorghum. Similarly, the four Arabidopsis ecotypes showed no consistent NPQ changes under drought. Two ecotypes for a seasonally dry and moderately wet region showed a significant increase in steady-state NPQ suggesting that upregulation of NPQ maximum level might play in these ecotypes an ecological adaptation (Fig. 5). The genotype-specific response of NPQ to drought was previously reported in maize, sorghum and rice (Baker et al., 2023; Dalal and Tripathy, 2018; Liu et al., 2012).

### Probing mechanism of NPQ regulation under stresses

The two main players in NPQ, PsbS and VDE, are sensitive to changes in luminal pH where low pH promotes PsbS protonation and VDE activity to enable NPQ (Li et al., 2004; Ruban and Wilson, 2021). The luminal pH can be regulated by two main processes, efflux of protons with the help of the chloroplast ATP synthase and influx of protons link to electron transport across the thylakoid membranes. To evaluate ATP synthase conductivity to protons (*g*H^+^) and relative changes in proton motive force (*pmf)*, we analyzed the ECSt parameter as the decay kinetics of the electrochromic shift (ECS) signal during brief dark intervals (Kuhlgert et al., 2016). The ECS signal reflects changes in the absorbance of chloroplast pigments in response to alteration in the electric field across the thylakoid membrane (Junge and Witt 1968; Witt 1979). The significantly higher NPQ, estimated from NPQ_max_, an asymptote of NPQ in light or/and NPQ_T_, only under chilling in soybean correlated with a significant increase in the thylakoid *pmf* (ECSt parameter; Fig. 2, Fig. 3B and F). As well, there was observed significant decrease in both *g*H^+^ and LEF (Fig. 3J and N). Slower ATP consumption under stress conditions reduces stroma inorganic phosphate availability and retards proton efflux via the ATP synthase (*g*H^+^), which would concomitantly increase *pmf* (Takizawa et al., 2007; Strand and Kramer, 2014; Shikanai and Yamamoto, 2017; Takagi et al., 2017). In addition to the release of H^+^ by oxygen-evolving complex of photosystem II, the Q-cycle in the cytochrome (Cyt) b6f complex contributes to the translocation of H^+^ to the lumen by the oxidation of plastoquinol. Increased acidification of the lumen (high ΔpH) slows the oxidation of plastoquinol at Cyt b6 f complex, thus regulating overall electron transfer between PSII and PSI (Kramer et al., 2004; Takizawa et al., 2007; Suorsa et al., 2012; Zia et al., 2016). Our results on chilling treated soybean suggest that the limitation in electron flow between PSII and PSI increased via pH-dependent restriction in plastoquinol oxidation at the cyt b6f complex (Takizawa et al. 2007). At the same time, the fraction of PSI center with oxidized P700 (P700^+^) under steady-state light did not change, suggesting that cyclic electron flow (CEF) could contribute to electron flow around PSI (Fig. 4B). The CEF would build up a high proton gradient across the thylakoid membranes to make then possible to drive further NPQ and ATP synthesis (Golding and Johnson, 2003; Shikanai and Yamamoto, 2017; Zia et al., 2016), with less generation of another energy source (NADPH) needed for Calvin-Benson cycle (CO_2_ fixation) (Kramer and Evans, 2011; Munekage et al., 2004; Rott et al., 2011; Yang et al., 2018).

Tobacco under both stresses, soybean and maize under drought stress showed an increase in NPQ despite no change in *pmf* and ATPase synthase activity (except soybean) compared to control plants (Fig. 3A-C, 3E-G, 3I-K). These results suggest that other mechanisms of NPQ upregulation could play a role here including accessibility of VDE coenzyme (Haupt and Glowacka, 2024), increase sensitivity of NPQ to ΔpH via stress accumulation of zeaxanthin (Zhang et al., 2009; Sacharz et al., 2017), activity of the xanthophyll cycle (Kubien and Sage, 2004), PsbS transcriptional regulation (Wei et al., 2019) as well as regulation of thylakoids ion channels and transporters (Spetea et al., 2017; Correa Galvis et al., 2020; Raju et al., 2024). Chloroplast *pmf* comprises both ΔpH and electrochemical gradient (ΔΨ), which contribute equally to ATP synthesis (Cruz et al., 2001) however only ΔpH contributes to the activation of NPQ. It was shown that changes in the partitioning of *pmf* into ΔpH and ΔΨ components can contribute to the regulation of NPQ (Kanazawa and Kramer 2002; Avenson et al. 2004; Kramer et al., 2004; Cruz et al., 2005; Takizawa et al., 2007). Shift towards prevail storage of *pmf* in ΔpH would make NPQ more sensitive to *pmf* changes and could be achieved by stress-induced regulation of thylakoid channels and transporters (Carraretto et al., 2013; Armbruster et al., 2014; Correa Galvis et al. 2020; Herdean et al., 2016). The shift in the partitioning of *pmf* towards more ΔpH would be particularly important in abiotic stresses in which the requirement for NPQ can increase, but LEF is too low to support a large enough *pmf* (Cruz et al., 2005). In addition, under drought treatment maize preserved similar *pmf* to this in control conditions despite the significant reduction of LEF (Fig. 3G and O), suggesting upregulation of CEF. A similar observation was made in two maize accessions with contrasting drought tolerance where the tolerant one could preserve CEF for a longer duration of stress than the susceptible one (Zhou et al., 2018). Under different stresses, NPQ response linked to change or stable *pmf*, chloroplastic-ATP synthase activity, and LEF have been reported (Kanazawa and Kramer, 2002; Kanazawa et al., 2017; Takagi et al., 2017; Yang et al., 2018; Raju et al., 2024).

We observed here a decrease in NPQ_T_ in sorghum (under both stresses) and in maize (under chilling) correlated with a *pmf* decrease, though both crops might utilize different mechanisms to achieve this phenotype (Fig. 3C, D and 3G, H). Under drought stress in sorghum, significantly lower *pmf* was achieved with significant upregulation of H^+^ conductivity of ATP synthase and stable LEF (Fig. 3H, 3L) with no significant changes in the redox state of PSI (Fig. 4D and 4H). These observations suggest no limitation on the PSI acceptor side which together with the mild reduction in stomatal conductance and C4-specific CO_2_ condensation mechanism, could suggest that the Calvin-Benson cycle acts as a substantial electron sink reducing the need for NPQ upregulation. Oppositely, under chilling maize significant reduction in NPQ that correlated with *pmf* decrease could result from a significant decrease in LEF (Fig. 3C, G, O). Since maize was shown to hold low chilling tolerance (Glowacka et al 2014), it might be speculated that chilling induced the impairment of PSII which would explain the decrease of both *pmf* and LEF (Hu et al 2023). The suppression of electron flow from PSII to PSI could then protect PSI from overreduction (no change in PSIor; Fig. 4G) and decrease the fraction of oxidized PSI centers under steady-state light (Fig. 4C). In the case of sorghum under the chilling, the significant reduction in NPQ was achieved without changes in *pmf* but with upregulation of LEF (Fig. 3 D, H, P). Since there was no decrease in the fraction P700 oxidized centers (P700^+^) under steady-state light (Fig. 4D), PSI centers could maintain their capacity to engage in phytochemistry. The significant increase in the fraction of the PSI centers that were over-reduced might suggest an increase limitation on the PSI acceptor side to pass the electrons (Fig. 4H).

### Lack of consistent NPQ phenotype across phenological stages under low-N stress

Under low-N, maize genotypes showed significant genetic variation in NPQ kinetics at seedling and pre-flowering stages (Fig. 6G and H; Fig. S5). NPQ induction rates in light and dark varied up to 1-fold between genotypes and developmental stages. NPQ phenotypes at the pre-flowering stage did not consistently align with those at the seedling stage under low-N stress. It was shown that maximum NPQ is significantly upregulated in the seedlings of rice cultivar with low sensitivity to low N, (Tantray et al., 2020) and that plant age influences the fate of excitation energy. The steady state of NPQ decreased with the age of the Arabidopsis plant possibly due to an increase in the photochemical quenching aligning with an increase in the Rubisco content (Johansson et al., 2004; Bielczynski et al., 2017). Oppositely, Carvalho et al. (2015) found that the maximum reversible photoprotective component of NPQ is highest during the reproductive phase, while juvenile and senescent phases exhibited lower values. Low soil fertility may intensify with the plant age, leading to less photochemical quenching and requiring increased NPQ with the plant transition from the seedling to pre-flowering stage. On the other hand, nitrogen depletion could lead to resource redistribution and photosystem II reorganization to absorb less energy which would create less need for NPQ. Genotype-dependent responses under low-N conditions reflect distinct expression patterns of NPQ-regulating genes related to chlorophyll catabolism like magnesium-dechelatase, light-harvesting chlorophyll *a*/*b*-binding protein, and N assimilation (Baker et al., 2023; Sahay et al 2024a, b). The interplay of this variety of mechanisms could explain the lack of uniform changes in NPQ across developmental stages under low-N stress observed here in maize.

## Conclusions

Here we demonstrated that leaf-disc fluorescence measurements are well suited to study NPQ across diverse species and stresses. This semi-high-throughput approach is revealed as a promising tool for the identification of NPQ kinetics regulatory genes in multiple crop species. Our examination of NPQ kinetics parameters under chilling, drought and low nitrogen led us to the conclusion of the existence of substantial variation in NPQ phenotypes across genotypes and environments. It appears that variations in NPQ kinetics resulting from genetic and environmental factors, are not separate from each other. Instead, different genotypes exhibit distinct plasticity of NPQ response to the same environmental perturbations. This underlines that studying only a single accession under stress conditions limits our understanding of the adaptation of whole species to excess light under stress conditions. We speculate that the lack of consistent NPQ phenotype across different stresses is due to specificities of some stresses, interplay of genetic variation in many NPQ regulatory genes and overlapping between short and long adaptations to excess light while others could be elucidated by differences between C3 and C4 type of photosynthesis. Across very species × stress combinations, a general trend of NPQ to follow the changes in *pmf* was not consistent. NPQ upregulated even when conductivity of protons in ATPase synthase and *pmf* were not changed as seen in tobacco under drought and chilling. However, the substantial increase by 5-fold in NPQ in soybean under chilling was related to change in *pmf* and a significant downregulation of chloroplast ATP synthase conductivity of protons. Based on the results we summarized the plausible mechanisms of NPQ modulation along with other NPQ-related parameters in four crops under chilling and drought conditions (Fig. 7). Our findings suggest that interpreting relative changes in NPQ phenotypes under abiotic stress is inherently complex and demands a broader integration of physiological data across multiple regulatory layers (Sahay et al., 2024b).

**Figure 7.**
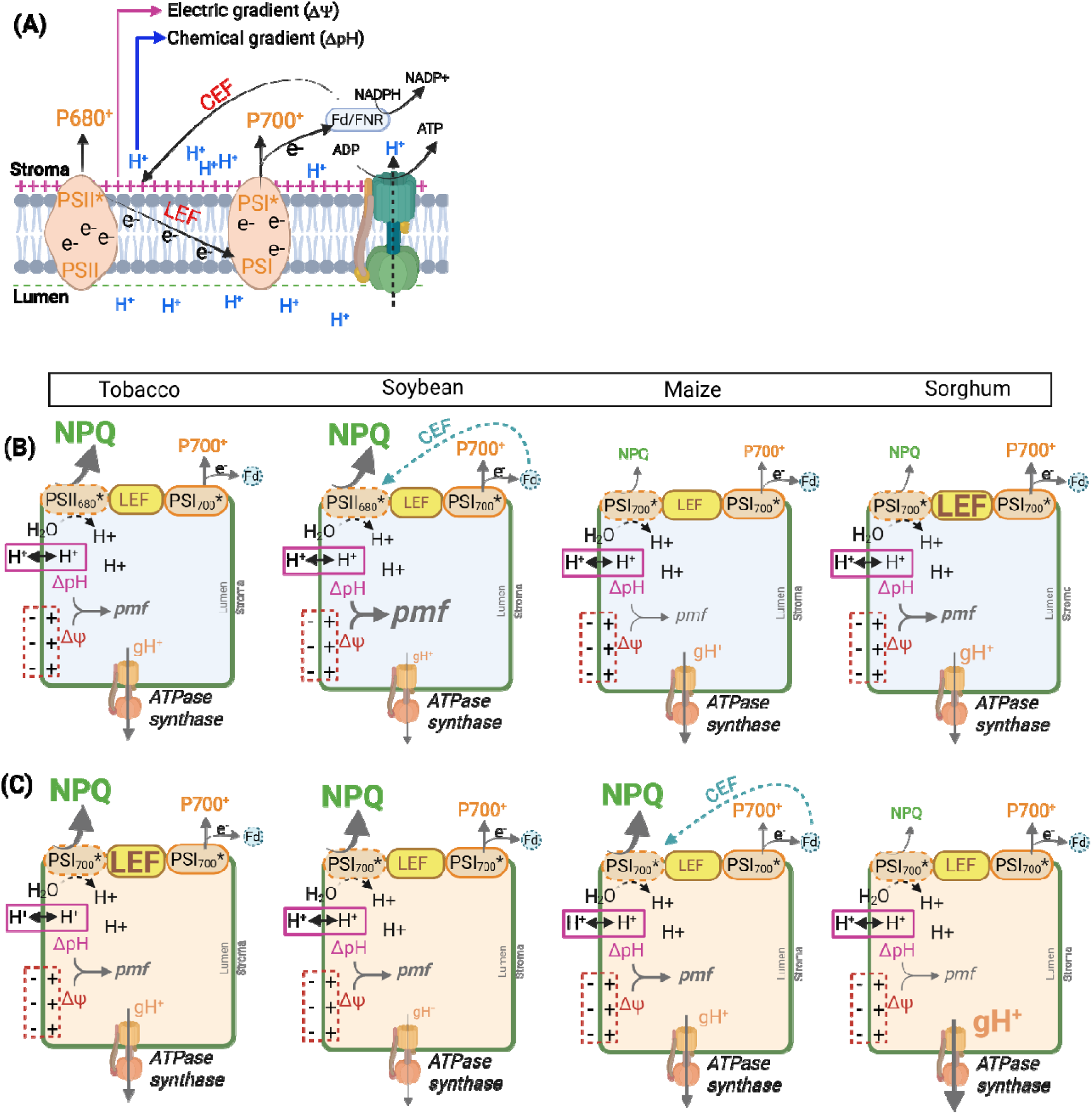
Plausible mechanisms underlying chilling- and drought-induced modulation of NPQ and associated parameters. **(A)** A general overview of photosynthetic mechanisms in plants. Linear electron flow (LEF) from photosystem II (PSII) to PSI leads to the accumulation of protons (H^+^) in the thylakoid lumen, generating a proton motive force (*pmf*) across the thylakoid membrane. The *pmf* consists of a H^+^ gradient (ΔpH) and an electrochemical gradient (ΔΨ), both of which are essential for driving ATP synthesis via chloroplast ATP synthase conductivity of protons (*g*H^+^). The ΔpH component of *pmf* plays a crucial role in regulating photosynthesis by reducing the efficiency of light capture through the energy-dependent quenching (qE) of NPQ mechanisms and slowing electron transfer to prevent damage to PSI reaction centers and reduce the production of harmful ROS in the thylakoid membrane. Proposed photoprotective strategies under **(B)** chilling and **(C)** drought stress conditions across four crops, tobacco, soybean, maize, and sorghum. Based on the results, NPQ modulation appears to be controlled by a combination of genetic and environmental factors. In panels B and C, thickness of the arrows and size of the fonts for *pmf*, *g*H^+^, LEF, NPQ, P700^+^ indicate relative change to control conditions in the corresponding parameter. Dotted lines represent parameters that were not directly measured in this study but are hypothesized to change as part of the proposed mechanism. CEF – cyclic electron flow; Fd- ferredoxin; P680* - excited state of chlorophyll in reaction center of PSII which is a primary donor of electron for PSI; P680^+^ - oxidized state of P680; P700* - excited state of chlorophyll in the reaction center of PSI. P700^+^ - oxidized state of P700. Figure was created using biorender.com and corresponds to Experiment II.

## Materials and Methods

### Experiment I: Comparison of different NPQ measurement approaches

To test the leaf-disc-(Method S1) and whole-plant-based approach to measure NPQ kinetics across plant species and stresses, we conducted trials on tobacco under chilling and soybean under drought stress. For tobacco, two transgenic lines psbs-4 (lower NPQ) and PSBS-28 (higher NPQ) and the wild type (WT) of *Nicotiana tabacum* cv. ‘Petite Havana’ (Glowacka et al., 2018) were used. Seeds were sown in 8.89 cm height × 8.89 cm width pots (SQN0350-0-B66, Hummert International, Earth City, MO, USA) filled with soil-less growing mix (10-12050, Berger BM2 germination mix; Hummert International). Pots were placed in a reach-in growing chamber (AR-66L2, Percival, Perry, IA, USA) set to a 16 h photoperiod, with temperature 18°C/21°C (night/day), 60% relative humidity, and a photosynthetic photon flux density (PPFD) of 200 µmol m^-2^ s^-1^. Trays (65-69630; Hummert International) holding the pots were covered with domes (15218200, Hummert International) to maintain moisture for germination by keeping a 2 cm water level. After 5-6 days, the domes were removed, and trays were repositioned every other day. Germinated seedlings of WT and transgenic lines were transplanted into trays (4 cm × 3 cm × 5.7 cm; 809 series, Hummert International) for two weeks and watered twice weekly with 150 ppm liquid fertilizer (Peter’s 20-10-20 general purpose fertilizer, 25#, Peters Inc., Allentown, PA, USA). To compare two NPQ measurement approaches, whole plant and leaf discs from the same plant, were imaged using a fluorescence imager. Initially, 2-weeks after germination whole plants were imaged and then placed back in the growth chamber for one to two days to restore normal physiology before leaf disc collection for NPQ assessment under control conditions. Concurrently, a separate set of WT plants was grown under identical growth-chamber conditions which were exposed to chilling stress at 4-6°C for 17 h (overnight chilling) by relocating them to a cold room. NPQ measurements were then conducted using both whole plant and leaf-disc-based approaches under chilling stress conditions as well.

To compare both the leaf-disc- and whole-plant-based approach to NPQ under different stress conditions in different plant species, wild type of soybean (*Glycine max* L. cv. ‘Throne’) was tested under PEG-6000 induced osmotic stress. Seeds were sterilized with 15% bleach for 7.5 minutes, thoroughly rinsed with sterile water, and then soaked overnight. The sterile seeds were germinated in a growth chamber (39940 Thermo-Kool, Mid-South Industries, Inc., Laurel, Mississippi) in the dark on a moist sterile Whatman 1 filter paper maintained at a PPFD 100 µmol m^-2^ s^-1^, with a temperature of 25°C and relative humidity of 55-60%. The germinated seedlings were transferred to hydroponic conditions, where they received either half-strength of autoclaved Hoagland media (pH 6.2-6.5; Methods S2) as the control or a stress treatment involving 10% polyethylene glycol 6000 (PEG 6000, Acros Organics, CAS, 25322-68-3), known as a drought inducer. The nutrient medium, with or without PEG, was regularly replaced to ensure adequate nutrition for the seedlings. Three weeks after germination, NPQ measurements were conducted on both whole plants and leaf discs from the same plants.

To compare the leaf-disc method that uses fluorescence imaging with multiphases flash method allowed with the use of fluorometer, the excised leaves in addition to leaf discs were collected from field-grown *S. bicolor* cv. ‘Tx430’. At the late-seedling stage, the youngest fully expanded leaf was excised at its base and submerged in a bucket containing water. The leaf was then recut under water just above the initial excision in the bucket, ensuring continuous submersion. Simultaneously, leaf discs were collected and placed into 96-well plates. Both plant materials were collected in the late afternoon, dark-adapted overnight at 4°C or room temperature, and subjected to fluorescence assays the following morning for NPQ quantification.

### Experiment II: NPQ in four crops across two stresses

We investigated the effect of chilling and drought stress on NPQ phenotypes across tobacco, soybean, maize, and sorghum by growing them side by side under growth-chamber conditions. The seeds of wild type of *Nicotiana tabacum* cv. ‘Petite Havana’, soybean (*Glycine max* L. cv ‘Throne’), maize (*Zea mays* L. cv. ‘B73’) and sorghum (*Sorghum bicolor* cv. ‘Tx430’) were grown in identical growth conditions as described for tobacco in above section except minor modification. Seeds were sown in identical pots (8.89 height × 8.89 width cm; SQN0350-0-B66, Hummert International, Earth City, MO, USA), and the germinated seedlings were thinned and kept one plant per pot. Seedlings received the same fertilizer regimen twice in a week, as described above. Three weeks after germination, plants were split into three groups: one group of plants was exposed to chilling stress at 4-6°C for 2 days by moving them to a cold room. The second group of plants was subjected to drought stress by withholding the watering for 5 days. The third set of the plants was maintained under regular conditions and used as control. NPQ measurements were conducted by imaging leaf discs from control, chilling, and drought stress conditions.

### Experiment III: NPQ across four Arabidopsis ecotypes under two abiotic stresses

The effect of both chilling and drought stress under fluctuating light on NPQ variation was studied in four *Arabidopsis thaliana* ecotypes originated from low and high elevation. Two low-elevation variants, CYR (52 m; France) and IP-Trs-0 (254 m; Iberia), as well as high-elevational variants, Sij-1 (2459 m; Uzbekistan) and Dja-1 (2995 m; Kyrgyzstan), of *Arabidopsis thaliana* were investigated for NPQ phenotypes under both chilling and drought stress. Seeds were stratified at 4°C for four days in the darkness. Subsequently, they were sown in soil-less potting mix (1220338; BM2 Germination and Propagation Mix; Berger, SaintModeste, Canada) in the same pots and a reach-in growth chamber described for tobacco (see first section of Material and Methods above). In this case, the growth-chamber maintained a temperature of 21°C during the day and 18°C at night, with an 8-hour light/16-hour dark photoperiod (200 µmol m^-2^ s^-1^) and 60% relative humidity. After one week of germination, seedlings were thinned to one per pot; plants were watered three times a week with 150 ppm liquid fertilizer (Peter’s 20-10-20 general purpose fertilizer, 25#, Peters Inc., Allentown, PA, USA). Additionally, plants were repositioned randomly within the chamber three times a week. After three-weeks, seedlings were split into two groups: one set of the seedlings was subjected to either drought conditions (with no watering for 5 days) or overnight chilling at 4-6°C, while the other set of seedlings remained under regular conditions and used as control. NPQ measurements were conducted by imaging whole plants under exposure to three cycles of high light intensity each followed by darkness. The leaf-disc approach was not used in Arabidopsis due to the soft leaf tissue, which makes it difficult to handle with forceps without causing substantial damage.

### Experiment IV: NPQ under low-nitrogen stress

NPQ response was investigated under low nitrogen conditions in maize (field) and teosinte (growth-chamber). Inbred maize (*Zea mays* cv. ‘B73’) and teosinte (*Zea mays* subsp. *parviglumis* cv. ‘Ames 21790’) were grown under low nitrogen (low N) stress in a growth-chamber (PGC20; Conviron Inc., Winnipeg, MB, Canada) controlled at 900 µmol m^-2^ s^-1^ PPFD, 70% humidity, a 16/8 h (maize) or 12/12 h (teosinte) light/dark photoperiod, and temperatures of 25/23°C day/night. The seeds were surface sterilized with 20% bleach for 20 minutes, thoroughly washed five times, and then soaked overnight in distilled water. Overnight soaked seeds were planted in 6-liter pots (13008000, Hummert International, Earth City, MO, USA) filled with growing mix (BM2 Germination and Propagation Mix; Berger, Saint-Modeste, Canada). Under control conditions, germinated seedlings were regularly supplied with full-strength Hoagland media. In contrast, for the low-N treatment, Ca(NO_3_)_2_ was completely replaced by CaCl_2_·2H_2_O, while KNO_3_ was reduced by 75% (for more details see Methods S2). NPQ measurements were conducted on leaf discs collected from 10 d (seedling) or/and 30 d (pre-flowering) old plants under control or/and low-N treatment conditions.

NPQ measurements were additionally carried out on 19 genotypes from the maize association panel grown under field conditions. The comprehensive details on the field experiment have been described by Sahay et al. (2024a and Methods S3). NPQ was analyzed in leaf discs collected from the youngest fully expanded leaves of 15-d and 31-d old plants after germination that corresponded to seedling and pre-flowering stages, respectively.

### Estimation of NPQ kinetics

The leaf discs were dark adapted for 17 hours before imaging, while whole plant imaging was performed after 20 minutes of dark adaptation periods consistent with the methodology described previously (Sahay et al., 2023). A schematic and detailed representation of the leaf disc-based approach of plant sampling and NPQ measurement is provided in Fig. S6 and Methods S1. Leaf discs, each with an area of 0.32 cm^2^, were collected using a hand-held puncher between 16:00 and 18:30 h. The discs were positioned with adaxial surface down into 96-well plates (781611; BrandTech Scientific, Essex, Ct, USA), covered with moist sponges, and incubated overnight in darkness. For the whole-plant NPQ measurement approach, the entire plant was subjected to a 20-minute dark incubation before imaging.

Both leaf discs and whole plant were imaged using a modulated chlorophyll fluorescence imager (FluorCam FC 800-C, Photon Systems Instruments, Drasov, Czech Republic). They were subjected to 10 min of either 1000 μmol m^-2^ s^-1^ light (for tobacco, and soybean) or 2000 μmol m^-2^ s^-1^ light (for sorghum, teosinte, and maize); comprising a combination of 500 or 1000 μmol m^-2^ s^-1^ of a red-orange light with λ_max_= 617 nm and 500 or 1000 μmol m^-2^ s^-1^ of a cool white 6500 K light), followed by 10 min of darkness (Experiments I, II and IV). The Arabidopsis plants were subjected to fluctuating light using three cycles of 3 min of 1000 μmol m^-2^ s^-1^ light each followed by 2 min of darkness (Experiment III). The minimum (*F*_o_) and maximal (*F_m_*) fluorescence in the dark were imaged. Subsequently, to capture changes in steady-state fluorescence (*F*_s_) and maximum fluorescence under illuminated conditions (*F*_m_’) over time, the saturating flashes of 3200 μmol m^-2^ s^-1^ (a cool white 6500 K light) in all other plant species except Arabidopsis for a duration of 800 ms were provided at the following intervals (in s): 15, 30, 30, 60, 60, 60, 60, 60, 60, 60, 60, 60, 60, 9, 15, 30, 60,180, and 300. For Arabidopsis, saturating flashes of 2400 μmol m^-2^ s^-1^ were used at the following intervals of 10, 10, 20, 20, 60, 60, 10, 20, 30, and 60s. NPQ at each time point was estimated using the Stern-Volmer quenching model (Bilger and Björkman, 1994) as described by equation 1:

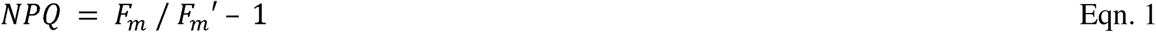

Maximum PSII efficiency (*F_v_*/*F_m_*) was estimated using the following equation 2 (Genty et al., 1989):

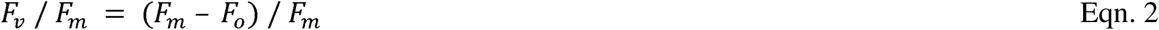

Raw images from leaf discs and whole plants were processed manually using FluorCam7. In leaf disc approach, individual leaf discs with an average area of 320 pixels were used in NPQ quantification. In the case of whole plant measurements NPQ quantification and analysis were performed on a leaf area of 170–180 pixels chosen manually from a portion of three of the youngest, fully expanded leaves subjected to the most uniform illumination.

The excised fully expanded leaves were incubated overnight in darkness before subjected to fluorescence assays the following morning. NPQ measurement was performed using an infrared gas analyzer LI-6800 (LiCOR, Lincoln, NE, USA) equipped with a fluorometer (6800-01A) using a multiphase flash routine (Loriaux et al., 2013). After the *F*_o_ and *F*_m_ fluorescence in the dark were recorded, the *F*_m_’ over a period of 10 minutes of illumination with 2000 μmol m^-2^ s^-1^ was collected using saturating flashes of 10,000 μmol m^-2^ s^-1^. The intervals between flashes were maintained similar to the program used to image leaf disc or whole plants in fluorescence imager.

### NPQ kinetics trait and data quality integrity

NPQ induction and relaxation values calculated at each time point were fitted to hyperbola and exponential equations in MATLAB (Matlab R2019b; MathWorks, Natick, MA, USA) to extract parameters reflecting the rate, amplitude, and steady state of the NPQ kinetics curve (Sahay et al. 2023). The complete list of the analyzed parameters and their biological meanings are presented in Table S1. Quality integrity was applied to data (Methods S4).

### Measurement of stomatal conductance

Stomatal conductance was measured on the tobacco, soybean, maize and sorghum under control, chilling and drought conditions (Experiment II) using the LI-600 (LiCOR) according to manufacture instruction.

### Measurements of other fluorescence- and absorbance- based photosynthetic parameters

Fluorescence- and absorbance-based parameters were measured using MultispeQ V2.0 (PhotosynQ, East Lansing, MI, USA; Kuhlgert et al., 2016). All the measurements were performed using Photosynthesis Rides protocol provided by the manufacturer, that is based on PAM method and data from MultispeQ were analyzed with the PhotosynQ web application (https://photosynq.org). Three-weeks old plants were measured across multiple crops under both chilling and drought conditions (Experiment II). A simplified NPQ parameter which does not require dark-adaptation (NPQ_T_), The thylakoid proton motive forces (*pmf*) *in vitro* (estimated from the ECSt parameter), thylakoid proton conductivity (gH^+^; an estimate of ATPase synthase), linear electron flow (LEF), PSI-oxidized centers, and PSI-over reduced centers were analyzed as a part of this study.

### Statistical analysis

Data were analyzed in SAS (v.9.4, SAS institute Inc., Cary, NC, USA). Data were tested with the Brown-Forsythe test for homogeneity of variance and the Shapiro-Wilk test for normality. In the cases when either of these tests failed the data were log-transformed. The significant effect of genotype, measurement approach, developmental stage, and treatment on absorbance and fluorescence delivered parameters were tested in ANOVA (α = 0.05). In Experiment II, significant effects in ANOVA were followed by a two-way test comparing the means of the treatments and that of the control (α = 0.05) for each species using Dunnett’s post-hoc test with multiple comparisons correction. In Experiment IV, the pair comparison between developmental stages inside the same accession for NPQ parameters were performed using t-test (α = 0.05). In Experiment I, a coefficient of determination for linear relation was used to determine a similarly between data delivered using leaf-disc and whole-plant measurements.

## Supplementary data

Supplementary figures, tables and methods described in this study are available in the online version of this article.

**Fig S1.** Comparison of leaf-disc and whole-plant based approach of NPQ measurement under control conditions.

**Fig S2.** Comparison of leaf-disc and whole-plant based approach of NPQ measurement under stress conditions.

**Fig S3.** Effect of fluctuating light in combination with chilling on NPQ kinetics in four *Arabidopsis thaliana* ecotypes.

**Fig S4.** Effect of fluctuating light in combination with drought stress on NPQ kinetics in four *Arabidopsis thaliana* ecotypes.

**Fig S5.** Effect of developmental stages on four NPQ traits in a set of 19 maize (*Zea mays*) genotypes.

**Fig S6.** Schematic representation of leaf-disc approach for NPQ measurement.

**Table S1.** List of NPQ kinetic traits with their description.

**Table S2** Comparison of maximum efficiency of PSII (*F_v_/F_m_*) between leaf-disc and whole-plant based measurement under control and stress conditions.

**Table S3** Absolute values of NPQ induction traits used for a comparative test between FlurCAM and LI-6800.

**Table S4** ANOVA table for NPQ traits in four *Arabidopsis thaliana* ecotypes in drought and chilling experiments.

**Table S5** ANOVA table for 6 NPQ traits in 19 maize (*Zea mays*) lines measured on seedling and pre-flowering stages.

**Method S1** Description of leaf-discs based sampling and NPQ measurement approach.

**Method S2** Description of Hoagland media-based low nitrogen and PEG treatment.

**Method S3** Corn field experimental design.

**Method S4** Data integrity and quality control procedure.

## Acknowledgments

The authors thank Annia Nelson, Bailey McLean, Rachel Gerdes, and Alexander Batelaan for assistant in growing and imaging of plant materials.

## Author contribution

SS performed the experiments, data analyses and graphing of results. DG and GB conducted the additional experiment and data analysis. KG, SS and JRL designed experiments and wrote the manuscript with input from GB and DG.

## Data availability

All data supporting the findings of this study are available within the paper and within its supplementary materials published online.

