## Supplementary material for "Nonphotochemical quenching changes with abiotic stressor and developmental stages": Supplementary Materials.docx

**Supplementary data**

Supplementary figures, tables and methods described in this study are available in the online version of this article.

**Fig S1.** Comparison of leaf-disc and whole-plant based approach of NPQ measurement under control conditions.

**Fig S2.** Comparison of leaf-disc and whole-plant based approach of NPQ measurement under stress conditions.

**Fig S3.** Effect of fluctuating light in combination with chilling on NPQ kinetics in four *Arabidopsis thaliana* ecotypes.

**Fig S4.** Effect of fluctuating light in combination with drought stress on NPQ kinetics in four *Arabidopsis thaliana* ecotypes.

**Fig S5.** Effect of developmental stages on four NPQ traits in a set of 19 maize (*Zea mays*) genotypes.

**Fig S6.** Schematic representation of leaf-disc approach for NPQ measurement.

**Table S1.** List of NPQ kinetic traits with their description.

**Table S2** Comparison of maximum efficiency of PSII (*F_v_/F_m_*) between leaf-disc and whole-plant based measurement under control and stress conditions.

**Table S3** Absolute values of NPQ induction traits used for a comparative test between FlurCAM and Licor6800.

**Table S4** ANOVA table for NPQ traits in four *Arabidopsis thaliana* ecotypes in drought and chilling experiments.

**Table S5** ANOVA table for 6 NPQ traits in 19 maize (*Zea mays*) lines measured on seedling and pre-flowering stages.

**Method S1** Description of leaf-discs based sampling and NPQ measurement approach.

**Method S2** Description of Hoagland media- based low nitrogen and PEG treatment.

**Method S3** Corn field experimental design.

**Method S4** Data integrity and quality control procedure.

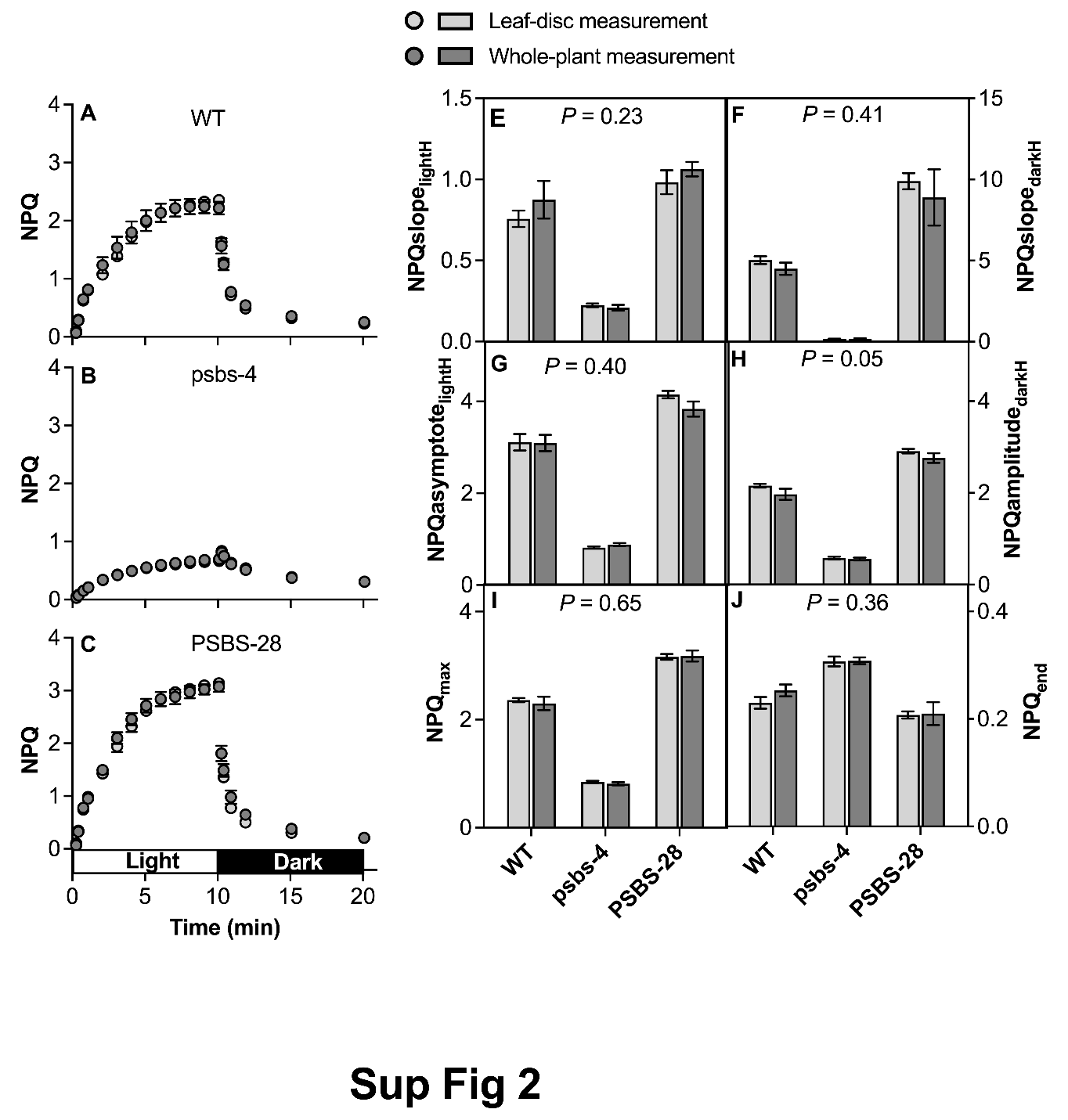

**Fig S1.** **Comparison of leaf-disc and whole-plant based approach of NPQ measurement under control conditions. A-C)** NPQ induction and relaxation kinetics measured in leaf discs and whole plant leaves of wildtype (WT) and two transgenic lines (psbs-4 silencing line and PSBS-28 overexpressing line) of tobacco (*Nicotiana tabacum*) plants during a 10 min light exposure (indicated by the white horizontal bar) followed by 10 min of dark exposure (indicated by the black horizontal bar). **D-I)** NPQ kinetics associated parameters derived from fitting hyperbolic (H) equation to NPQ curves in the light and dark. Traits: **E-F)** NPQslope_lightH_ and NPQslope_darkH_ describe the rate of NPQ in the light and dark, respectively. **G)** NPQasymptotelightH describes the steady-state of NPQ in the light. **H)** NPQamplitudedark describes the range of NPQ relaxation. **I-J)**. NPQmax and NPQend are the last values of NPQ in light and dark, respectively. A full definition of each trait is provided in Table S1). Data represent the mean ± SEM (n = 8 biological replicates, each from two to three technical replicates). The *P*-value indicates the effect of the measuring approaches on the kinetics traits in ANOVA. Data present results of Experiment I.

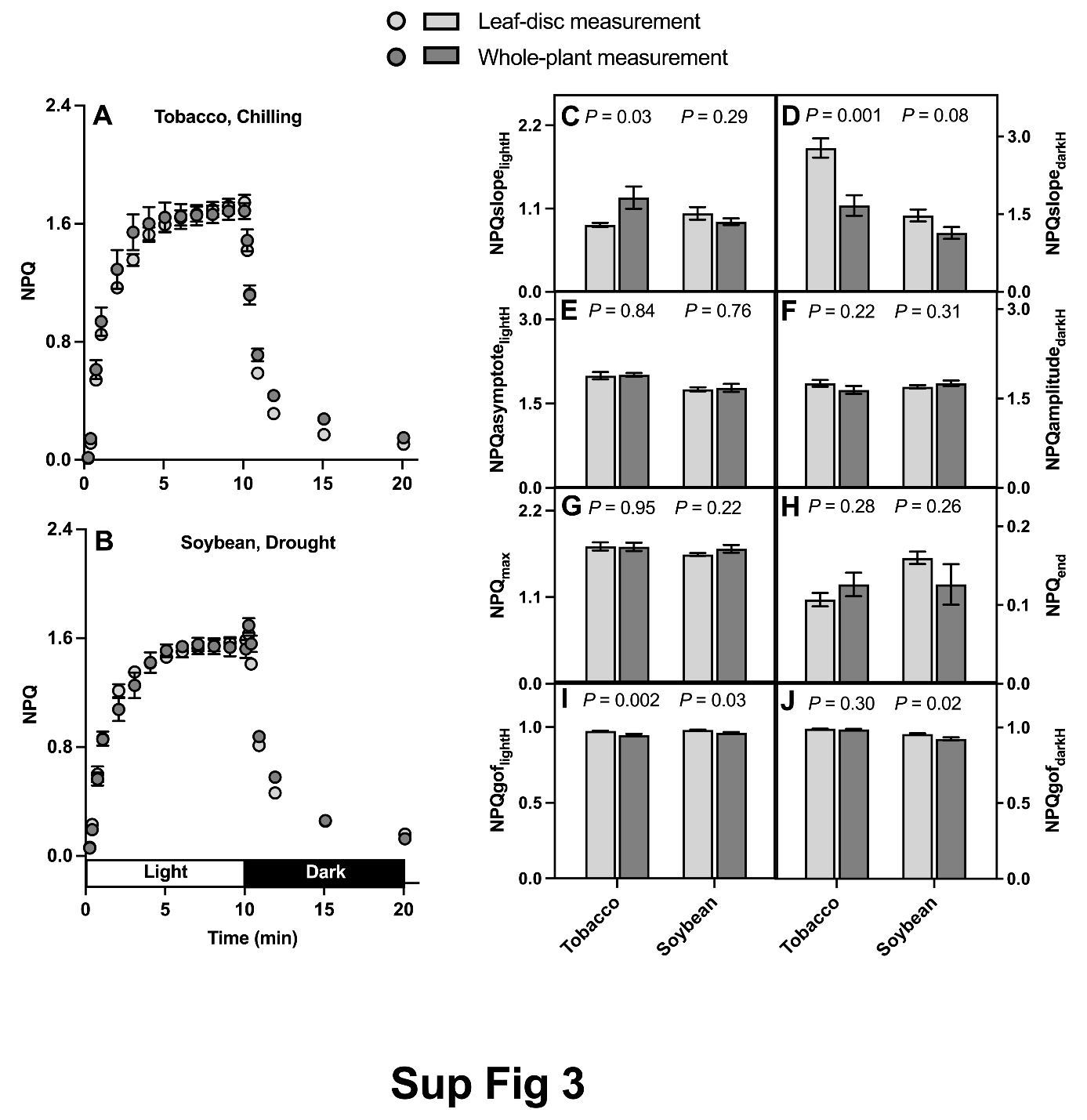

**Fig S2.** **Comparison of leaf-disc and whole-plant based approach of NPQ measurement under stress conditions.** **A-B)** Kinetics of NPQ induction and relaxation measured in leaf disc and whole plant leaves of tobacco (*Nicotiana tobacum*) and soybean (*Glycine max*) plants grown under chilling and drought stress conditions, respectively. **C-J)** NPQ kinetics associated parameters derived from fitting hyperbolic (H) equation to NPQ curves in the light and dark. Traits: **C, D)** NPQslope_lightH_ and NPQslope_darkH_ describe the rate of NPQ in the light and dark, respectively. **E)** NPQasymptote_lightH_ describes the steady-state of NPQ in the light. **F)** NPQamplitude_dark_ describes the range of NPQ relaxation. **G, H)** NPQ_max_ and NPQ_end_ are the last value of NPQ in light and dark, respectively. **I, J)** NPQgof_lightH_ and NPQgof_darkH_ describe the goodness of fit to the curve of NPQ in the light and dark, respectively. A full definition of each trait is provided in Table S1). Data represent the mean ± SEM (tobacco, n = 8; soybean, n = 4-5 biological replicates, each from two to three technical replicates). The *P*-value indicates the effect of the measuring approach on the kinetics traits in ANOVA. Data present results of Experiment I.

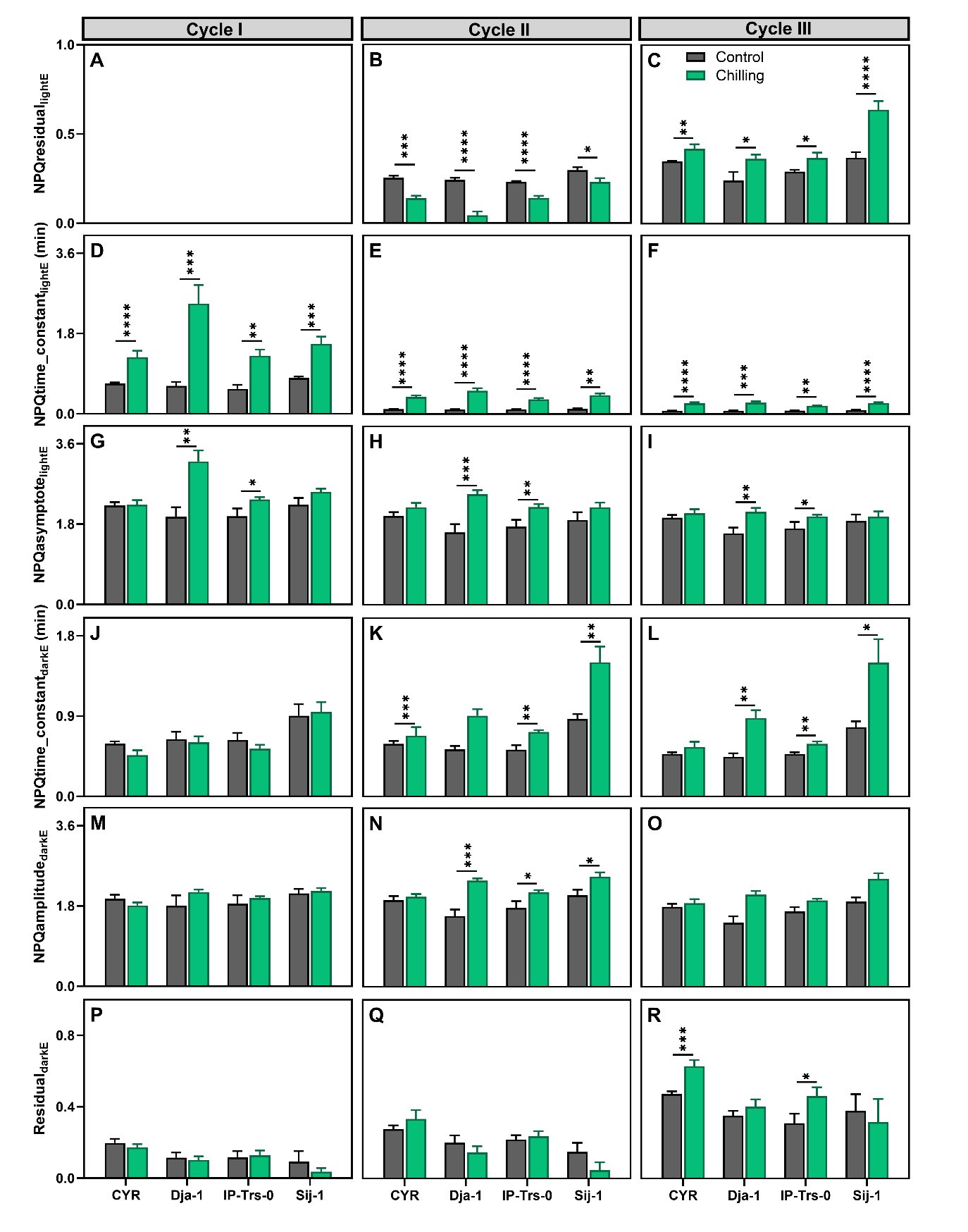

**Fig S3. Effect of fluctuating light in combination with chilling on NPQ kinetics in four *Arabidopsis thaliana* ecotypes.** Kinetics traits of NPQ induction and relaxation in chilling stress under three subsequent light-dark cycles in four ecotypes represent different elevations (in the bracket): CYR (52 m), Dja-1 (2995 m), IP-Trs-0 (254 m), and Sij-1 (2459 m). Traits: (A-C) NPQresidual_lightE_ describes the initial NPQ prior to light exposure, (D-F) NPQtime_constant_lightE_ describes the time period for NPQ induction following light exposure (min), (G-I) NPQasymptote_lightE_ describes the steady-state NPQ in the light, (J-L) NPQtime_constant_darkE_ describes the time period for NPQ induction in the dark (min), (M-O) NPQamplitude_darkE_ describes the range of NPQ relaxation; and (P-R) NPQresidual_darkE_ describes the remaining NPQ (residual) after dark exposure. A full definition of each trait is provided in Table S1. Each trait is estimated for three sequential light-dark cycles. Data represent means ± SEM (n = 5-6 biological replicates). Asterisks indicate significant differences in the t-test between chilling and corresponding control values (**P* < 0.05; ***P* < 0.01; ****P* < 0.001, and *****P* < 0.0001). Data present results of Experiment III.

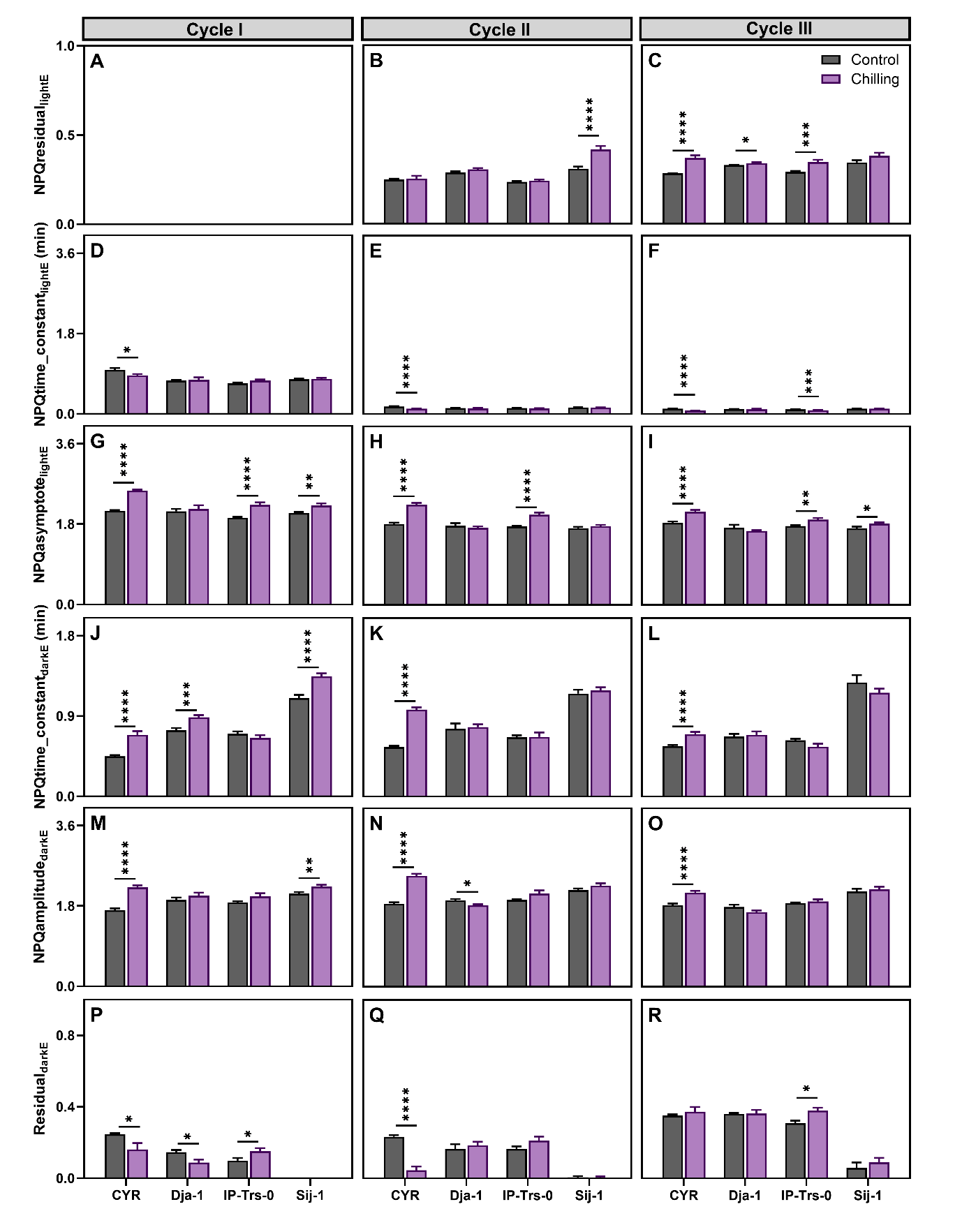

**Fig S4.** **Effect of fluctuating light in combination with drought stress on NPQ kinetics in four *Arabidopsis thaliana* ecotypes.** Kinetics traits of NPQ induction and relaxation in drought stress under three subsequent light-dark cycles in four ecotypes represent different elevations (in the bracket): CYR (52 m), Dja-1 (2995 m), IP-Trs-0 (254 m), and Sij-1 (2459 m). Traits: (**A-C**) NPQresidual_lightE_ describes the initial NPQ prior to light exposure, (**D-F**) NPQtime_constant_lightE_ describes the time period for NPQ induction following light exposure (min), (**G-I**) NPQasymptote_lightE_ describes the steady-state NPQ in the light, (**J-L**) NPQtime_constant_darkE_ describes the time period for NPQ relaxation in the dark (min), (**M-O)** NPQamplitude_darkE_ describes the range of NPQ relaxation; and (**P-R**) NPQresidual_darkE_ describes the remaining NPQ (residual) after dark exposure. A full definition of each trait is provided in Table S1. Each trait is estimated for three sequential light-dark cycles. Data represent means ± SEM (n = 5 (only Dja-1) to 8 biological replicates). Asterisks indicate significant differences in the t-test between drought and corresponding control values (**P* < 0.05; ***P* < 0.01; ****P* < 0.001, and *****P* < 0.0001). Data present results of Experiment III.

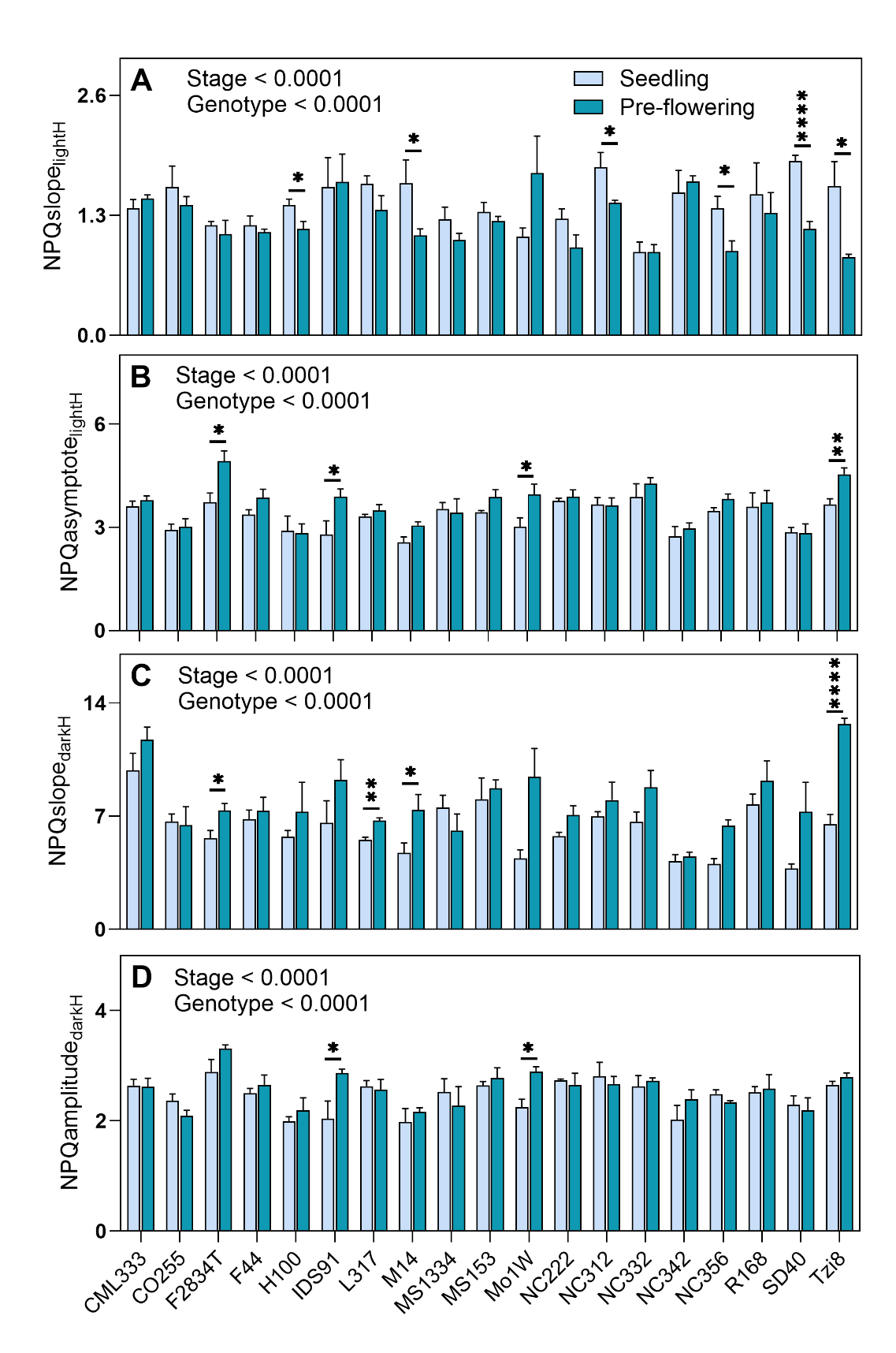

**Fig S5.** **Effect of developmental stages on four NPQ traits in a set of 19 maize (*Zea mays*) genotypes.** NPQ kinetics was measured at seedlings and pre-flowering stages in a set of 19 maize genotypes grown under low-N field conditions. **(A)** NPQslope_lightH_ and NPQslope_darkH_ describe the rate of NPQ in the light and dark, respectively. NPQasymptote_lightH_ describes the steady-state of NPQ in the light. NPQamplitude_darkH_ describes the range of NPQ relaxation. The NPQ_max_ and NPQ_end_ are the last values of NPQ in light and dark, respectively which are presented in Fig. 6G, H in the main manuscript. A full definition of each trait is provided in Table S1**.** Data represent means ± SEM (n = 4 biological replicates). Asterisks indicate significant differences in the t-test between corresponding developmental stages (**P* < 0.05; ***P* < 0.01; ****P* < 0.001, and *****P* < 0.0001). Data present results of Experiment IV.

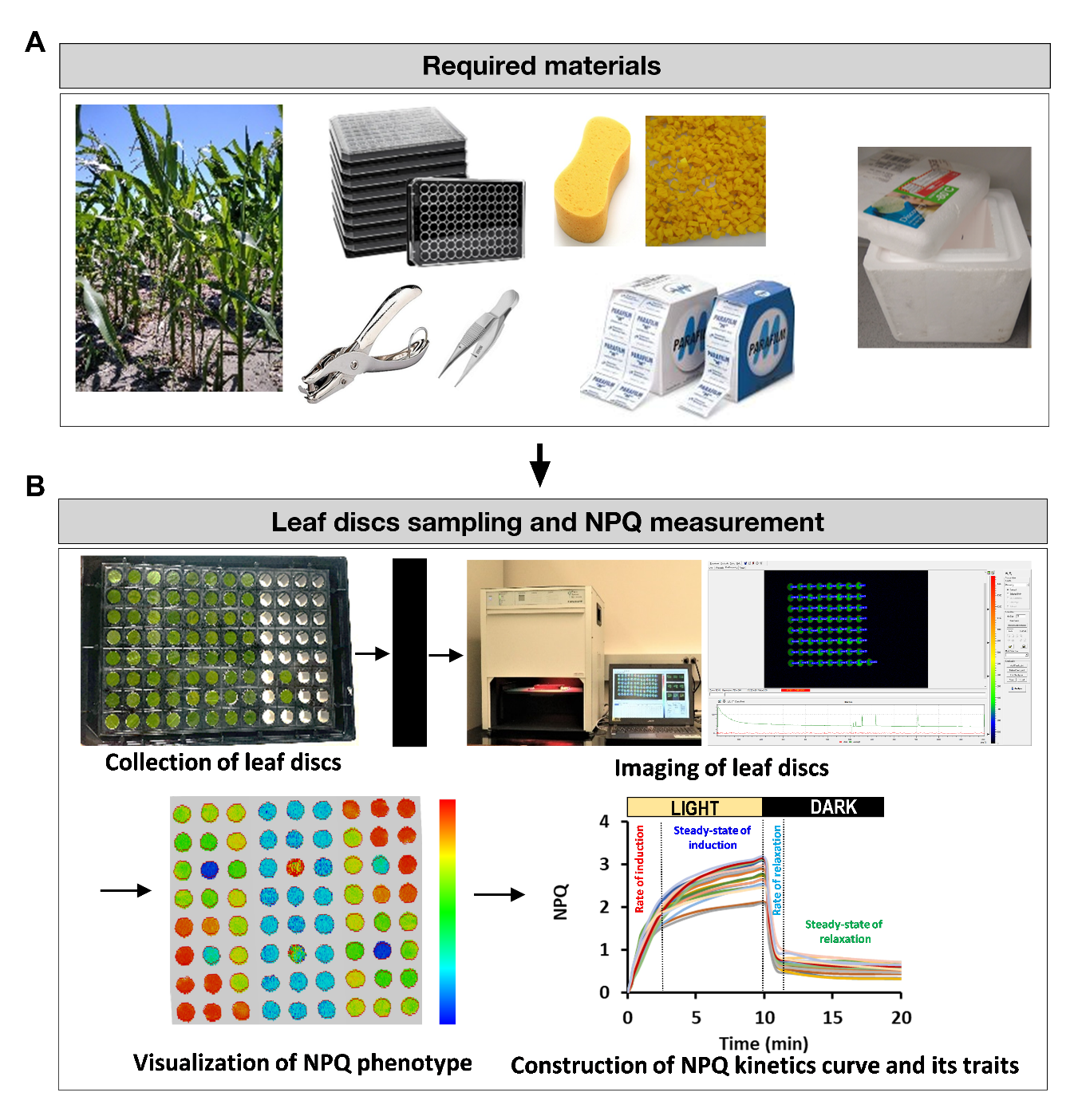

**Fig. S6** **Schematic representation of leaf-disc approach for NPQ measurement**. **(A)** Required materials needed for the sampling of leaf discs from the plants. **(B)** A flow diagram showing a collection of leaf discs into 96-well plate, dark incubation, imaging, and analysis of measured and type of the derived parameters.

**Table S1** List of NPQ kinetic traits with their description.

| **Kinetics Type** | **Kinetics Attributes** | **Trait Abbreviation Name** | **Trait Full Name** | **Trait Description** |
| --- | --- | --- | --- | --- |
| NPQ induction in light | Rate | *NPQslope_lightH_* | Initial slope of NPQ in light | How fast NPQ is induced under light. Estimated from a hyperbolic (H) curve fit to the different measurements of NPQ collected during the 10-minute light treatment. |
|  |  | *NPQtime_constant_lightE_* | Time constant of NPQ in light | Estimate of the time at which NPQ reaches 63% of its final maximum potential value after being exposed to high light (min), calculated from exponential (E) function. |
|  | Range | *NPQamplitude_lightH_*  *NPQamplitude_lightE_* | NPQ amplitude in light | Size of the change between estimated maximum NPQ induction and pre-induction NPQ, calculated from hyperbolic (H) and exponential (E) fit. |
|  | Steady state | *NPQasymptote_lightH_*  *NPQasymptote_lightE_* | Asymptote of NPQ in light | The maximum potential value NPQ reaches under prolonged light. Estimated from a hyperbolic (H) and exponential (E) function. |
|  |  | *NPQ_max_* | Maximum NPQ | The highest of NPQ during the light treatment. |
| NPQ relaxation in light | Rate | *NPQslope_darkH_* | Initial slope of NPQ in dark | How fast NPQ relaxes once the lights are turned off. Estimated from a curve fit to a hyperbolic (H) function. |
|  |  | *NPQtime_constant_lightE_* | Time constant of NPQ in dark | Estimate of the time at which NPQ reaches 63% of its final maximum potential value after being exposed to darkness (min), calculated from exponential (E) function. |
|  | Range | *NPQamplitude_darkH_*  *NPQamplitude_darkE_* | NPQ amplitude in dark | Size of the change between estimated maximum NPQ induction and minimum NPQ during relaxation, calculated from hyperbolic (H) and exponential (E) fit. |
|  | Steady state | *NPQresidual_darkE_* | Residual of NPQ in dark | The amount of potential NPQ relaxation beyond that which occurs during the dark treatment, calculated from exponential (E) function. |
|  |  | *NPQ_end_* | Minimum NPQ in dark | Remaining NPQ still present at the end in the dark. |

**Table S2** **Comparison of maximum efficiency of PSII (*F_v_/F_m_*) between leaf-disc and whole-plant based measurement under control and stress conditions.** *F_v_/F_m_* was measured in control and chilling conditions in wildtype, psbs-4 (silencing line), PSBS-28 (overexpressing line) tobacco (*Nicotiana tabacum*) plants, and under drought conditions in wildtype soybean (*Glycine max*) plants. Data represents the means ± SEM (tobacco, n = 8; soybean, n = 4 biological replicates, each from two to three technical replicates). The *P*-value indicates the effect of the measurement approach ANOVA. Data present results of Experiment I.

| Method | Tobacco | | | Soybean |
| --- | --- | --- | --- | --- |
|  | Wildtype | psbs-4 | PSBS-28 |  |
|  | Control | | |  |
| Leaf-disc measurement | 0.81 ± 0.004 | 0.81 ± 0.003 | 0.82 ± 0.001 | - |
| Whole-plant measurement | 0.83 ± 0.001 | 0.83 ± 0.001 | 0.82 ± 0.002 | - |
| *P-*values | 0.07 | | |  |
|  | Chilling | | | Drought |
| Leaf-disc measurement | 0.78 ± 0.002 | 0.78 ± 0.002 | 0.78 ± 0.004 | 0.81 ± 0.002 |
| Whole-plant measurement | 0.77 ± 0.006 | 0.77 ± 0.007 | 0.77 ± 0.005 | 0.79 ± 0.001 |
| *P-*values | 0.05 | | | 0.11 |

**Table S3 Absolute values of NPQ** **induction traits used for a comparative test between FlurCAM and Licor6800.** Measurements were taken on control and chilling treated sorghum (*Sorghum bicolor*). Data correspond to relative changes showed in Fig 1F-H in the main manuscript and represents the means ± SEM. Data present results of Experiment I.

| NPQ Parameter | FluorCAM | | LiCOR | |
| --- | --- | --- | --- | --- |
|  | Control | Chilling | Control | Chilling |
| NPQslope_lightH_ | 1.23 ± 0.07 | 1.78 ± 0.13 | 0.65 ± 0.10 | 1.03 ± 0.09 |
| NPQaymptote_lightH_ | 2.72 ± 0.091 | 2.32 ± 0.07 | 2.49 ± 0.11 | 2.14 ± 0.11 |
| NPQ_max_ | 2.48 ± 0.08 | 2.33 ± 0.08 | 1.98 ± 0.07 | 1.91 ± 0.08 |

**Table S4 ANOVA table for NPQ traits in three subsequent light-dark cycles for four *Arabidopsis thaliana* ecotypes in drought and chilling treatments.** The *P*-values are bolded for traits that significant effect of factor and/or factor interaction was detected. Data present here correspond to Fig. 5 in the main manuscript and Fig. S3 and Fig. S4 in the Supplementary data file. Data present results of Experiment III.

| NPQ traits | *P-*value | | | | | |
| --- | --- | --- | --- | --- | --- | --- |
|  | Drought treatment | | | Chilling treatment | | |
|  | Treatment (T) | Ecotype (E) | T x E | T | E | T x E |
|  | *Light-dark cycle I* | | | | | |
| NPQasymptote_Light_ | **<.0001** | **<.0001** | **0.0006** | **<.0001** | 0.20 | **0.01** |
| NPQtime constant_Light_ | 0.88 | **<.0001** | **0.06** | **<.0001** | **0.009** | **0.02** |
| NPQamplitude_Dark_ | **<.0001** | **<.0001** | **<.0001** | 0.49 | 0.25 | 0.42 |
| NPQrate constant_Dark_ | **<.0001** | **<.0001** | **<.0001** | 0.28 | **0.0008** | 0.49 |
| NPQresidual_Dark_ | 0.08 | **<.0001** | **0.005** | 0.31 | **0.008** | 0.73 |
|  | *Light-dark cycle II* | | | | | |
| NPQasymptote_Light_ | **<.0001** | **<.0001** | **0.0003** | **<.0001** | 0.79 | **0.05** |
| NPQtime constant_Light_ | **0.0004** | **0.0029** | **0.0002** | **<.0001** | 0.32 | **0.01** |
| NPQresidual_Light_ | **0.02** | **<.0001** | 0.08 | **<.0001** | **<.0001** | 0.10 |
| NPQamplitude_Dark_ | **<.0001** | **<.0001** | **<.0001** | **<.0001** | **0.01** | **0.01** |
| NPQrate constant_Dark_ | **<.0001** | **<.0001** | **<.0001** | **<.0001** | **<.0001** | 0.08 |
| NPQresidual_Dark_ | **0.02** | **<.0001** | **<.0001** | 0.66 | **<.0001** | 0.13 |
|  | *Light-dark cycle III* | | | | | |
| NPQasymptote_Light_ | **<.0001** | **<.0001** | **0.004** | **0.002** | 0.49 | 0.23 |
| NPQtime constant_Light_ | **<.0001** | **<.0001** | **<.0001** | **<.0001** | 0.38 | 0.06 |
| NPQresidual_Light_ | **<.0001** | **0.0006** | **0.02** | **<.0001** | **<.0001** | 0.23 |
| NPQamplitude_Dark_ | **0.02** | **<.0001** | **0.002** | **<.0001** | **0.001** | **0.04** |
| NPQrate constant_Dark_ | 0.45 | **<.0001** | **<.0001** | **<.0001** | **<.0001** | **0.02** |
| NPQresidual_Dark_ | **0.01** | **<.0001** | 0.15 | **0.04** | **0.005** | 0.52 |

**Table S5 ANOVA table for 6 NPQ traits in 19 maize (*Zea mays*) lines measured on seedling and pre-flowering stages.** Plants were grown under low nitrogen field conditions. The *P*-values are bolded for traits that significant effect of factor and/or factor interaction was detected. Data present here correspond to Fig. 6G-H in the main manuscript and Fig. S5 in the Supplementary data file. Data present results of Experiment IV.

| NPQ traits | *P-*value | | |
| --- | --- | --- | --- |
|  | Stage (S) | Genotype (G) | S x G |
| NPQslope_lightH_ | **<.0001** | **<.0001** | **0.01** |
| NPQaymptote_lightH_ | **<.0001** | **<.0001** | 0.25 |
| NPQ_max_ | **0.001** | **<.0001** | 0.29 |
| NPQslope_darkH_ | **<.0001** | **<.0001** | **0.01** |
| NPQaymptote_darkH_ | 0.10 | **<.0001** | 0.33 |
| NPQ_end_ | 0.12 | **<.0001** | 0.12 |

**Method S1** **Description of leaf-discs based sampling and NPQ measurement approach.**

**Required materials**

1. Black 96-well plate with coverlid (Thermo Scientific cat no 167008)
2. Pieces of sponge
3. Single-hole puncher (¼ inch circular hole, item #825232, Office Depot Inc.)
4. Tweezer/Forceps
5. Aluminum foil
6. Parafilm
7. Styrofoam box with ice packs, which are taped to the coverlid
8. PAM-fluorescence imager (e.g. Closed FluorCam FC 800-C)

**Preparatory steps for sampling the leaf discs from growth chamber or field setups**

1. Before sampling, ensure all necessary materials are prepared before sampling.
2. Punch the leaf disc in an order of experimental set up using hole puncher and place the dorsal (top) side of leaf disc facing down onto the bottom of plate using tweezer.
3. Once the leaf disc is positioned upside down, place sponge pieces behind the leaf disc to cover it.
4. After completing the collection of leaf discs, cover the plate with a lid and wrap it with aluminum foil.
5. Store the plates with the transparent bottom surface facing up in a Styrofoam box and incubate them in the dark overnight (17 hours) at room temperature to minimization of the effect of microenvironmental conditions at the time of sampling like light intensity, temperature, and time of day and allow the relaxation of qE and qZ.
6. Following day, unwrap the plate from the aluminum foil and image it using a fluorescence imager (Closed FluorCam FC 800-C, Photon Systems Instruments, Drasov, Czech Republic) with a custom-created script for 10 minutes of induction and 10 minutes of relaxation in FluorCam 7 software.
7. Analyze the raw images by excluding the background, save average pixel values within the area in numeric form (.TXT), and export the data to a spreadsheet. Plot NPQ induction and relaxation curves against 13 light and 6 dark time-points. In the case of whole-plant, process the raw image manually by drawing a circle of known area of the uniformly light-exposed fully expanded leaves.
8. Fit the response of NPQ to time to the hyperbola and exponential equation to obtain the other kinetics-associated parameters using custom-made script in MATLAB software.

**Important tips and recommendations for leaf disc sampling**

1. **Photosynthetic Rate Variation:** It is important to note that the dorsal surface of the leaf has a different photosynthetic rate than the ventral surface. Collect leaf discs from the middle part, slightly away from the midrib, of the youngest fully developed leaf. This ensures consistency, as the flag leaf (uppermost) or fully matured (lowermost) leaf may exhibit entirely different photosynthetic characteristics than the youngest fully developed leaf.
2. **Proper Plate Handling:** Hold the plate using your thumb and index finger to prevent interference with the bottom transparent surface. Avoid placing the plate on your palm or soil while collecting leaf discs into wells, as it may compromise image and data quality. Using a coverlid attached to the plate with tape offers a better option to rest the plate or keep the bottom surface away from potential interference.
3. **Optimal Moisture Level:** Ensure that sponge pieces are slightly moist, avoiding excessive moisture that could affect leaf discs and induce hypoxia conditions. If water seeps from the sponge while pressing it between finders, the sponge is too wet.
4. **Precise Positioning:** It is necessary to verify that the leaf discs are well-positioned against the bottom of the 96-well plate. If not, make adjustments by gently moving them to the center of the wells.
5. **Inverted Plate Storage:** Store plates overnight in an inverted position (bottom side up). This allows any excessive water from the wells, including sponge pieces, to drip out, if present.
6. **Temperature Considerations:** In hot summer field conditions, use a cool Styrofoam box with a lid and an ice pack to store plates. This prevents leaf discs from drying out due to high temperatures. Ensure plates do not directly contact ice packs to avoid freezing leaf discs. In the fall, a Styrofoam box without an ice pack can be used.
7. **Dark Room Imaging:** The process of unwrapping and imaging the plate in the dark room should be done to maintain dark adaptation of plant material.
8. **Uniform Light Exposure:** For avoiding the ununiform light exposure during the fluorescence assay place the plate in the center of the imager tray.

**Material S2** **Description of Hoagland media- based low nitrogen and PEG treatment.**

**Hoagland Media (Arnon, 1956):** Hoagland media is one of the commonly utilized mediums for cultivating plants in both hydroponic and soil conditions. Soybean plants were grown in full strength of Hoagland media as control and 10% polyethylene glycol (prepared in Hoagland medium) as drought stress inducer in hydroponic conditions. Teosinte plants were grown in low nitrogen conditions by applying modified concentrations of nitrate salts in Hoagland medium into the soil. The composition details for the full-strength (2M) Hoagland medium, encompassing stock and working concentrations, are provided below:

| **Hoagland Full nutrient solution (1L)** | | | | | |
| --- | --- | --- | --- | --- | --- |
| Chemicals | Amount (g/L) | Stock solution  (250 mL) | Working solution  (mL/ L) | Stock Concentration | Final working concentration |
| **Macronutrients** | | | | | |
| KNO_3_ | 202.2 | 50.55 | 2.5 | 2M | 5 mM |
| Ca(NO_3_)_2_ 4H_2_O | 472.3 | 118.075 | 2.5 | 2M | 5 mM |
| KH_2_PO_4_ (pH 6.0) | 136.09 | 34.023 | 1 | 1M | 1 mM |
| MgSO_4_ 7H_2_O | 492.94 | 123.323 | 1 | 2M | 2 mM |
| **Micronutrients** | | | | | |
| H_3_BO_3_ | 2.86 | 0.715 | 1 | -- | 0.046M |
| MnCl_2_ 4H_2_O | 1.81 | 0.465 |  | -- | 0.009M |
| ZnSO_4_ 7H_2_O | 0.22 | 0.055 |  | -- | 7.5×10^-4^M |
| CuSO_4_ | 0.08 | 0.02 |  | -- | 3.2×10^-4^M |
| H_2_MoO_4_ 4H_2_O | 0.02 | 0.005 |  | -- | 1.11×10^-4^M |
| **Iron source** |  |  |  |  |  |
| 0.05M | Fe-EDTA* | 2.8 | 0.7 | 1 | 50 µM |
| *3.72 g Na-EDTA in 70 mL ddH_2_O mixed with 2.78 g FeSO4.7H_2_O in 70 mL ddH_2_O and then volume up to 200 mL. | | | | | |
| **Hoagland – Low N nutrient solution (1L)** | | | | | |
| **Macronutrients** | | | | | |
| KNO_3_ | 50.55 | 12.63 | 2.5 | 0.5M | 1.25 mM |
| CaCl_2_ 2H_2_O | 294.02 | 73.505 | 2.5 | 2M | 5 mM |
| KH_2_PO_4_ (pH 6.0) | 136.09 | 34.023 | 1 | 1M | 1 mM |
| MgSO_4_ 7H_2_O | 492.94 | 123.323 | 1 | 2M | 2 mM |
| **Micronutrients*** | -- | -- | 1 | -- | -- |
| *Same as given above | | | | | |
| **Iron source** | Fe-EDTA* | 2.8 | 0.7 | 1 | 50 µM |

**Half strength Hoagland solution**: For half-strength Hoagland media, dilute the full nutrient solution by 50% using autoclave water.

**PEG 6000 (10%) working solution**: Prepare 10% PEG solution by dissolving 100 g in 1 L of autoclaved Hoagland media.

**Method S3 Corn field experimental design.**

Each genotype was replicated across four blocks under low nitrogen treatment conditions. The low nitrogen experimental plots were arranged with two rows of the same genotypes, spaced ~75 cm apart, and individual plants within the same row were approximately 15 cm apart. Each plot extended to a length of 1.5 meters, with planting of 11 kernels per row.

**Method S4** **Data integrity and quality control procedure.**

Data extracted from both leaf discs and whole plant measurements were subjected to a quality control and outlier removal procedure based on *F*_v_/*F*_m_ and goodness of fit for hyperbolic and exponential equations fit to the measured NPQ. In control treatment conditions, *F*_v_/*F*_m_ values 0.58 were used as cut-off and any values lower than 0.58 were excluded. However, in abiotic stress treatments, *F*_v_/*F*_m_ cut-off was not applied, as the effects of abiotic stresses on *F*_v_/*F*_m_ were considered. The goodness of fit was defined as the discrepancy between measured values and the values predicted by the equation fit to the data. Measurements with poor fits (below 1^st^ percentile) were excluded for both control and stress treatment conditions. After quality control, data for NPQ was used for statistical analysis.
